# Improve consensus partitioning via a hierarchical procedure

**DOI:** 10.1101/2021.09.03.458844

**Authors:** Zuguang Gu, Daniel Hübschmann

## Abstract

Consensus partitioning is an unsupervised method widely used in high throughput data analysis for revealing subgroups and assigns stability for the classification. However, standard consensus partitioning procedures are weak to identify large numbers of stable subgroups. There are two main issues. 1. Subgroups with small differences are difficult to separate if they are simultaneously detected with subgroups with large differences. And 2. stability of classification generally decreases as the number of subgroups increases. In this work, we proposed a new strategy to solve these two issues by applying consensus partitionings in a hierarchical procedure. We demonstrated hierarchical consensus partitioning can be efficient to reveal more subgroups. We also tested the performance of hierarchical consensus partitioning on revealing a great number of subgroups with a DNA methylation dataset. The hierarchical consensus partitioning is implemented in the R package *cola* with comprehensive functionality for analysis and visualizations. It can also automate the analysis only with a minimum of two lines of code, which generates a detailed HTML report containing the complete analysis.

## Introduction

Consensus partitioning or consensus clustering is an unsupervised method that classifies samples into subgroups and evaluates the stability of the classification by resampling from original data (Monti et al.). It has become an important tool applied in high-throughput data analysis, *e*.*g*., to reveal cancer subtypes (Sturm et al. 2012) or to validate the agreement of the classification to known clinical factors. In our previous work (Gu et al. 2020), we developed an R/Bioconductor package named *cola* that provides a general framework for consensus partitioning. It allows simultaneously running multiple feature-selection methods and partitioning methods and it provides comprehensive visualization and reporting utilities for automatic and deep comparison and interpretation on the results. *cola* provides a new and efficient method named *ATC* (ability to correlate to other rows) for extracting top features and it recommends spherical *k*-means clustering (Hornik et al. 2012) for subgroup classification. Through comprehensive benchmarks on public datasets, we demonstrated *cola* was able to generate new, stable and biologically meaningful classifications.

*cola* provides a convenient toolkit for performing consensus partitioning analysis. It performs well when the expected number of subgroups is relatively small, *e*.*g*., no larger than 6 as demonstrated in (Gu et al. 2020). However, when the number of expected subgroups increases, issues for general consensus partitioning procedures (Wilkerson and Hayes 2010; Kiselev et al. 2017) rise and they would significantly affect the classification. In consensus partitioning procedures, top *n* features scored by a certain method, *e*.*g*. standard deviation, are firstly selected. Later, sample classification is only applied on the top features. A good classification expects the selected features to have the ability to separate all subgroups, in other words, consensus partitioning procedures take into account all samples equally. However, in real-world datasets, this condition cannot always be met. It is possible that features good at separating major subgroups (*i*.*e*. subgroups with large difference) are weak for secondary subgroups (*i*.*e*. subgroups with small difference) if the secondary subgroups have different sets of features that are efficient for classification. When the real number of subgroups becomes larger, it is highly possible that subgroups have different sets of efficient features for classification, and this leads to secondary subgroups being difficult to gain stable separation if classifying them with major subgroups at the same time. The second issue is when the number of subgroups gets larger, the probability of two samples not in the same subgroup tends to increase, which results in the loss of stability of the classification. Both issues restrict the classification to reach a large number of stable subgroups.

In this work, to solve the previously mentioned issues, we propose a strategy named *hierarchical consensus partitioning* (HCP) that applies standard *cola* consensus partitioning (CP) in a hierarchical procedure. Simply speaking, one could first classify samples into *k* groups where *k* is a small number which corresponds to major subgroups. Then for each subgroup of samples, one could repeatedly apply consensus partitioning with a new set of top features extracted only to that subset of samples. The hierarchical procedure stops until certain criteria are reached. By this means, theoretically, small subgroups or secondary subgroups could be detected in later steps of the hierarchical procedure. This process can generate a hierarchy of subgroups where subsets of samples are represented as nodes in the hierarchy. The idea of executing consensus partitioning hierarchically has also been applied in identifying consensus network modules to reveal multiresolution modularity of the network (Jeub et al. 2018).

For large datasets with huge numbers of samples, in early steps of the hierarchical procedure, numbers of samples in the subsets could still be large. Due to that consensus partitioning by-nature is a time-consuming analysis, to improve the efficiency of partitioning on large datasets, we propose a strategy which randomly picks samples to a small subset, on which consensus partitioning is only applied, later the class labels of the unselected samples are predicted based on the classification of the selected samples. This down-sampling strategy ensures analysis of thousands of samples can be done in an acceptable time period.

HCP extended the *cola* framework and it has been implemented in the *cola* package from version 2.0.0. For submatrices represented as nodes in the subgroup hierarchy, standard CP by *cola* is applied where specific combinations of feature selection methods and partitioning methods can be either user-defined or be selected from the built-in methods. HCP provides rich visualizations for interpretation of the results, as well as comprehensive tools for downstream analysis, such as dimension reduction, signatures analysis and functional enrichment analysis if signatures can be corresponded to genes. For the ease of use, HCP automates the analysis only with a minimum of two lines of code, which generates a detailed HTML report containing the complete analysis.

In the paper, we first illustrated the issues of CP with a simulated dataset and a real-world dataset. Then we demonstrated the use of HCP with an RNA-sequencing (RNAseq) dataset with an intermediate number of samples. The results showed HCP was able to reveal more subgroups compared to standard CP analysis. Next we applied HCP on a single-cell RNA-sequencing (scRNAseq) dataset with a large number of cells, where the down-sampling functionality was enabled in the analysis. The results showed HCP classification was similar to the one from the original study but cell clusters had larger separation under HCP classification. Last, we tested the performance of HCP on revealing a great number of subgroups with a DNA methylation dataset.

## Methods

### A brief introduction to consensus partitioning and the *cola* package

The HCP method proposed in this work is an extension of the consensus partitioning implemented in the R package *cola*. To make the paper easy to read and self-explorary, here we briefly describe the methods and terms used in *cola*. Readers please refer to the original publication for more details (Gu et al. 2020).

Consensus partitioning is applied on columns of matrix-like data to discover subgroups of samples. Matrix rows are firstly assigned with scores by a certain method such as the widely used standard deviation (SD), then top *n* features with the highest scores are only used for consensus partitioning. This is called the *top-value method* in *cola*. We proposed a new top-value method *ATC* (ability to correlate to other rows) in *cola* which aims to capture the top features that are potentially highly correlated to other features and to provide more consistent patterns for subgroup classification. For row *i* in a matrix, denote the variable *X* as a vector of absolute values of the correlation coefficients to all other rows, the ATC score for row *i* is defined as:

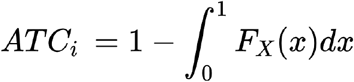

where *F*_*X*_(*x*) is the cumulative distribution function (CDF) of *X*. The aim of using top *n* features for partitioning is to keep the informative features that help partitioning while removing other features with irrelevant noise. We demonstrated that the ATC can capture better and distinct features for partitioning that cannot be captured by other methods (Gu et al. 2020).

After top *n* features are selected, a certain partitioning method is repeatedly applied on a randomly sampled subsets (*e*.*g*. 80%) of features and the stability of the partitioning is evaluated from the list of individual partitioning results, *i*.*e*., how often two samples stay in the same subgroup. According to extensive benchmarks performed in our previous study (Gu et al. 2020), we demonstrated that the spherical *k*-means clustering (skmeans) could classify samples more efficiently with higher stability and it also revealed more subgroups.

To get the optimized number of subgroups in consensus partitioning, a list of numbers of subgroups (denoted as *k*) are tried. The best *k* is evaluated by several metrics. *cola* mainly uses three metrics to determine the best *k*: mean silhouette score, PAC score and concordance scores, which measure the stabilities of the consensus partitioning from different aspects. Silhouette score measures how close one sample is to its own subgroup compared to the closest neighbouring subgroup and the mean of silhouette scores over all samples are used to measure the overall stability of the classification; PAC scores measures the proportion of ambiguous clusterings (Şenbabaoğlu et al. 2014) where the probability of two samples in the same group is between 0.1 and 0.9 (The form of 1-PAC is actually used in *cola* to let the direction of changes be the same as the other two metrics); And concordance scores measures the agreement of individual partitions to the consensus partition.

The *cola* package implements a comprehensive framework and also an easy interface for consensus partitioning analysis. It allows various user-defined methods easily integrated into different steps of the analysis, *e*.*g*., for feature selection, sample classification or definition of signatures. *cola* provides a complete set of tools for comprehensive subgroup analysis, including partitioning, signature analysis, functional enrichment, as well as rich visualizations for interpretation of the results. Moreover, to find the method that best explains a user’s dataset, cola allows running multiple methods simultaneously and provides functionalities for a straightforward comparison of results.

### Consensus partitioning methods are weak to simultaneously distinguish major and secondary subgroups

We demonstrated this issue with a simulated dataset. Two random matrices denoted as **M**_1_ and **M**_2_ were generated with 100 columns and 20 columns respectively. Both matrices had 100 rows. **M**_1_ was simulated as a set of samples from a two-condition comparison where the first 50 columns were labelled as group “A1” and the second 50 columns were labelled as group “A2”. Rows in the two groups were assigned with different levels of difference. For row *i* in **M**_1_, values in group A1 were generated from a normal distribution *N*(*μ*_1,*i*_, 1) and values in group A2 were generated from *N*(-*μ*_1,*i*_, 1). To simulate that rows in **M**_1_ had varied differences in the two groups, the vector of mean values ***μ***_1_ was generated from the standard normal distribution *N*(0, 1), thus, standard deviation of row *i* in **M**_1_ increases when *μ*_1,*i*_ has a higher absolute value. Similarly, **M**_2_ was also simulated as a set of samples from a two-condition comparison where the first 10 columns were labeled as “B1” and the second 10 columns were labelled as “B2”. To simulate **M**_2_ as a matrix with smaller row differences, rows in **M**_2_ were generated from *N*(0.5*μ*_2,*i*_, 1) and *N*(−0.5*μ*_2,*i*_, 1) where the vector of mean values ***μ***_2_ was also generated from *N*(0, 1). The order of values in ***μ***_2_ was set in a way that rank of the absolute values of ***μ***_2_ is identical to the reverse rank of the absolute values of ***μ***_1_, *i*.*e*., *rank*(|***μ***_2_|) ≡ *rank*(-|***μ***_1_|)). In this setting, if row *i* shows the highest difference between group A1 and A2, it shows the smallest difference between group B1 and B2.

**M**_1_ and **M**_2_ were merged into a single matrix denoted as **M** where A1/A2 were groups with major differences in **M** and B1/B2 showed relatively smaller differences. Figure 1A illustrates the heatmap for the random dataset. According to the column dendrogram on the heatmap, the four groups were located in the four separated branches. The separation of the four groups was also confirmed by the principle component analysis (PCA) in Figure 1B, but the columns were separated into three groups in the first principle component which explained 46% of the total variance of **M** while B1 and B2 were only separated in the second principle component which only explained 3% of the total variance. CP performed with *cola* was applied on **M** where top 50 rows with the highest standard deviation were selected as features and *k*-means clustering was applied to classify samples. The CP result showed that the best number of subgroups was three according to the empirical cumulative density function (eCDF) curve of the consensus values (the probability of two samples in a same subgroup) where a horizontal line extended almost from 0 to 1 for *k* = 3 (Figure 1C). This means in the three-group classification, in the repetitive classifications by resampling from the complete feature set, any two-sample pair was either always in the same group (consensus value close to 1) or belonged to different groups (consensus value close to 0). This can also be confirmed by the membership heatmap which visualized every single partitioning result where the samples in all 50 partitionings almost had the same classifications (Figure 1D). But, when the number of groups was set to four, the membership heatmap showed, in approximately 90% of the individual partitionings, either group1 or group2 was further split into two smaller groups, while only in 10% of all partitionings, B1 and B2 were correctly separated. This misclassification resulted in CP being hard to assign B1 and B2 as stable classifications (Figure 1E), thus it rejected four as the optimized number of subgroups. This problem is mainly due to that almost all the top 50 rows with the highest SD from **M** also had the highest SD in **M**_1_, thus efficient to separate A1 and A2; while these 50 rows had very small SD if only counting **M**_2_, which resulted that, these features were not good at separating B1 and B2 (Figure 1F).

**Figure 1.**
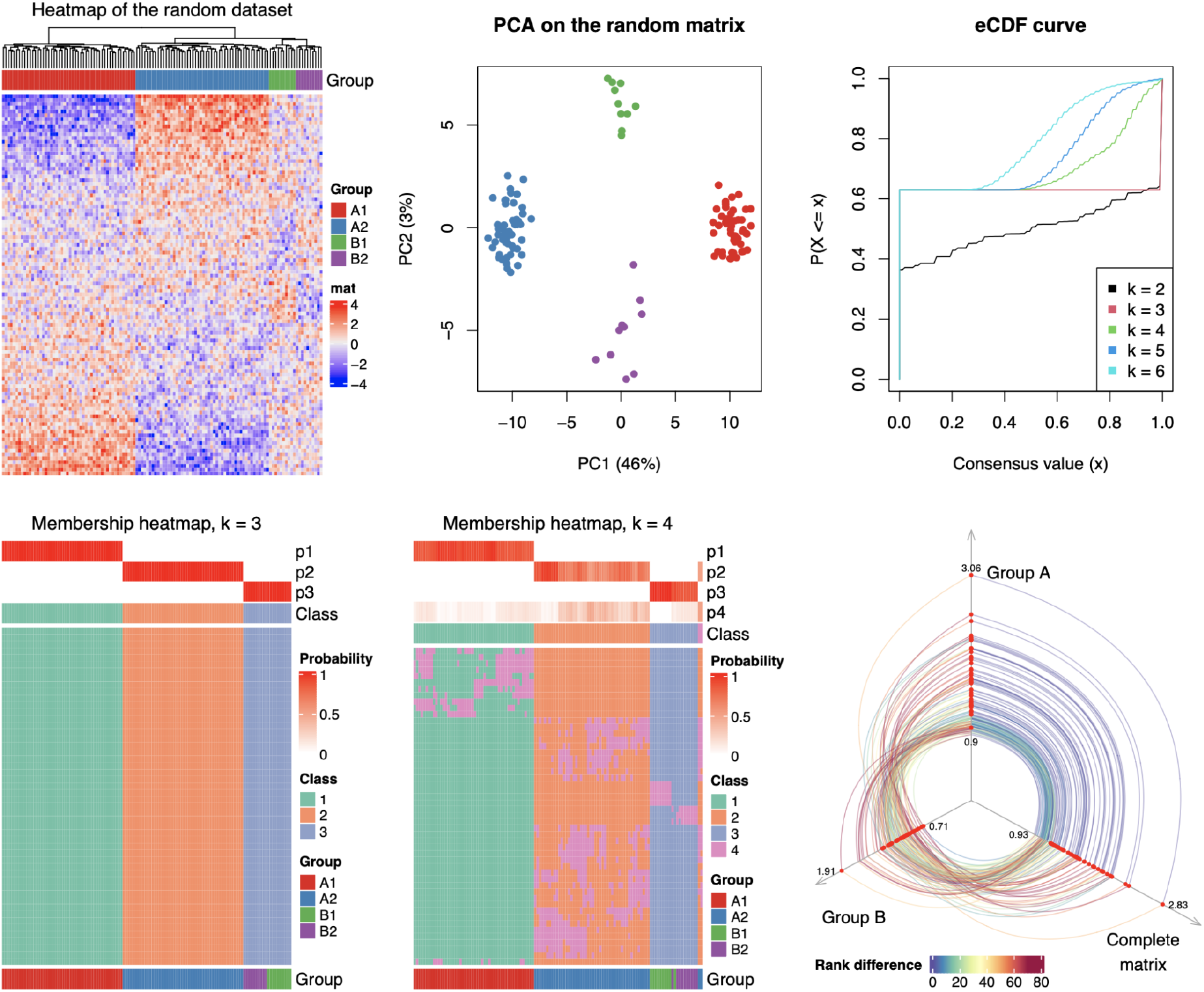
Consensus partitioning methods are weak to simultaneously distinguish major and secondary subgroups. A) Heatmap of the simulated dataset. B) Principal component analysis of the simulated dataset. C) Empirical CDF curves of the consensus partitioning for each *k*. D) Membership heatmap of the consensus partitioning with *k* = 3. E) Membership heatmap of the consensus partitioning with *k* = 4. In D and E, top annotations with names p1∼p4 correspond to the probability of samples belonging to each subgroup. F) Compare top 50 features in different groups. The 50 features are with the highest SD calculated from the complete simulated matrix. The three axes correspond to SD values calculated in matrices of group A, group B and the complete matrix. The top 50 features are highlighted in red dots in the three axes. The same features are connected between axes and the lines are colored by the rank difference of the SD values in the two corresponding matrices. The numbers on the three axes represent ranges of SD values from the three matrices.

Here we demonstrated a simulated dataset only with four groups. In the real-world dataset, if the expected number of subgroups is large, it is highly possible that the hierarchical structure exists and secondary subgroups would be hidden if applying standard CP to all samples.

Thus a method to capture the hierarchical structure of data is needed. In Supplementary File 1, we demonstrated HCP is able to detect all of the four groups of this random dataset.

### Consensus partitioning are less stable for larger *k*

The second issue for standard CP procedures is when the expected number of subgroups increases, the probability of two samples that are not in the same subgroup tends to increase as well, which results in the decrease of the stability of the classification for larger *k*. We demonstrate this issue with the human skeletal muscle myoblasts (HSMM) scRNAseq dataset (Trapnell 2021), on which we applied CP with ATC as top-value method, skmeans as partitioning method and number of subgroups were tried from 2 to 8. The heatmap of the top 1000 genes with the highest ATC scores suggested there should be a large number of subgroups (*e*.*g*. > 10) with clear patterns (Figure 2A). However, the optimized number of subgroups selected by CP can only reach a small value. Figure 2B∼2E illustrate the selection of the best *k* by the eCDF curves and three metrics of 1-PAC score, mean silhouette score and concordance score. Consensus partitionings showed high stability with *k* between 2 and 6 where the scores of 1-PAC scores, mean silhouette scores and concordance scores were very close to 1, although mean silhouette scores and concordance scores decreased slightly when *k* increased. When *k* exceeded 6, scores of 1-PAC, mean silhouette and concordance dropped dramatically compared to previous *k*. If the selection of the best *k* is based on the maximal votes from the three metrics with their highest values, *k* = 2 was taken as the best result, and if robustness is allowed, *k* = 6 might be the best result. Nevertheless, the two *k* (2 and 6) were still far from the expected one, which cannot be assigned with high stability under the framework of CP. The decrease of stability can also be observed in Figure 2F which directly visualized the membership of every individual partition (membership heatmap) and the probability of every two samples in the same subgroup (consensus heatmap). It showed when *k* becomes larger (e.g. *k* = 7 and 8), samples tended to show ambiguous classifications in individual partitionings.

**Figure 2.**
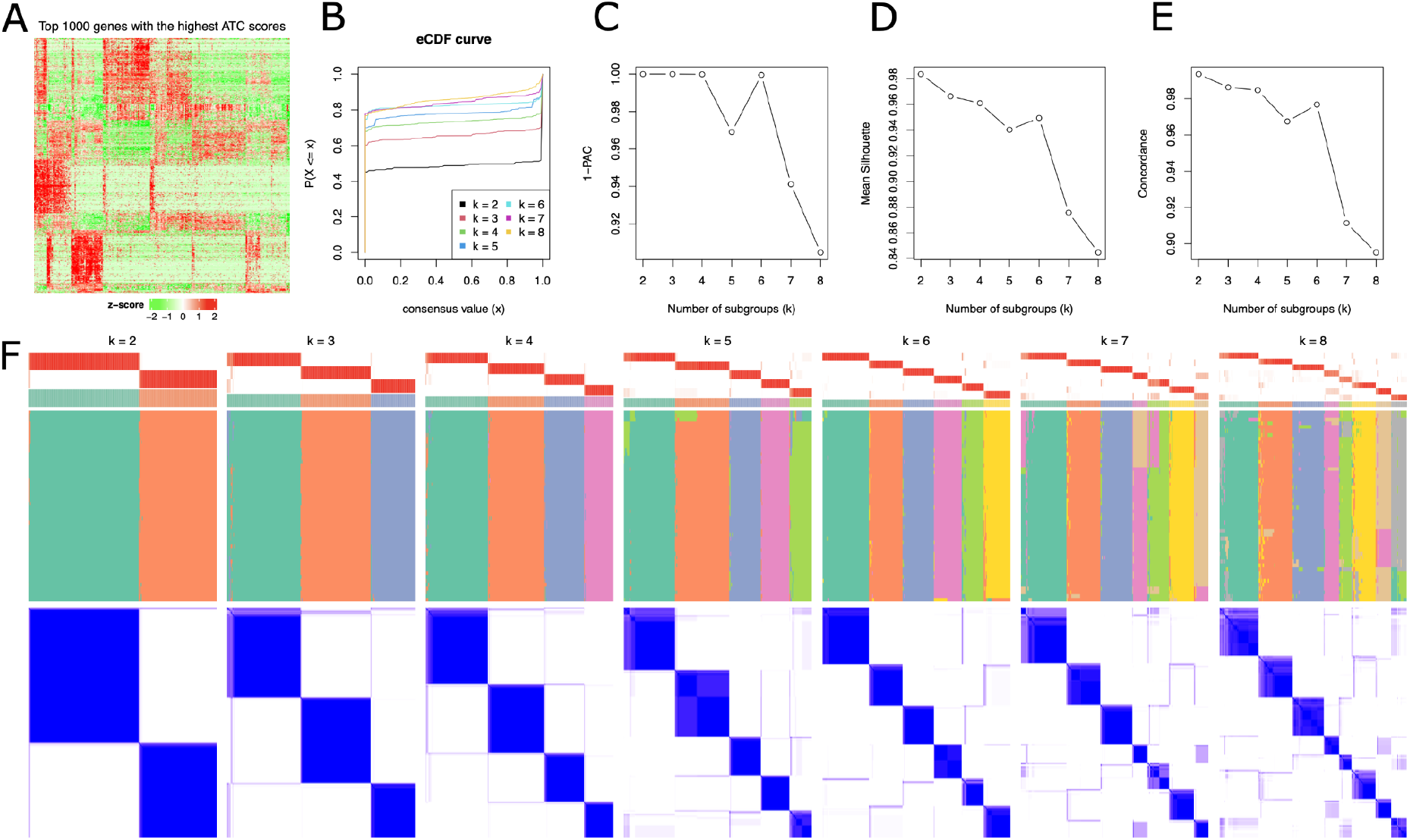
Illustration of the issue of large *k* in consensus partitioning with the HSMM scRNAseq dataset. A) Heatmap of top 1000 genes with the highest ATC scores. B) eCDF curves. C)1-PAC scores verse *k*. D) mean silhouette scores verse *k*. E) Concordance score verse *k*. F) An integrated visualization of the clustering results. Each column corresponds to the results of a specific *k*. For each *k*, from the top to bottom, there are the following plots: 1. An annotation showing the probability of samples in each subgroup; 2. The consensus classification; 3. Membership heatmap that visualizes all individual partitionings; 4. The consensus heatmap that visualizes the probability of every two samples in the same subgroup. For each *k*, columns have the same orders for all plots.

This example implies CP has a preference to assign high stability to small *k*, while it is difficult for large *k* to be stable. This issue can also be solved by applying CP with a hierarchical procedure where on each step of the iteration, CP is only applied with small *k* that ensures the gain of stability and more subgroups can be found in later steps of the hierarchical procedure.

### Workflow of the hierarchical consensus partitioning procedure

We propose a method named *hierarchical consensus partitioning* (HCP) to perform consensus partitioning via a hierarchical procedure. The HCP procedure is illustrated in Figure 3. The complete matrix is taken as the input of the whole analysis and the full set of samples are taken as the root node labelled as “0” in the hierarchy. CP with a single combination of a top-value method and a partitioning method or multiple combinations of both methods are applied on the matrix with a list of *k* (Figure 3, step 1) and the best *k* from the best method is selected (Figure 3, step 2). If there are multiple methods showing stable partitionings, the one with the highest number of signatures is selected. Note *k* should be set up to a small number. After the best partitioning is selected, next there are two filters to decide whether the samples should be split according to the classification. First the stability measured by mean silhouette score is tested against a cutoff (Figure 3, step 3). If the best partitioning is not stable, the whole set of samples are treated as unclassible and the node is treated as a leaf in the hierarchy. If it is stable, next whether the separation has biological meanings is tested by the number of signatures in the classification (Figure 3, step 4). If the number of signatures is sufficient, The partitioning result on the node is accepted and subgroups are taken as its child nodes (Figure 3, step 5). For each child node, we then test whether there exist sufficient numbers of samples of the corresponding submatrix. HCP is iteratively applied to each child node if there are enough samples (Figure 3, step 6). In HCP, these filterings can also be applied when the subgroup hierarchy is completely generated and users can manually fine-tune the hierarchy by merging or further splitting the nodes.

**Figure 3.**
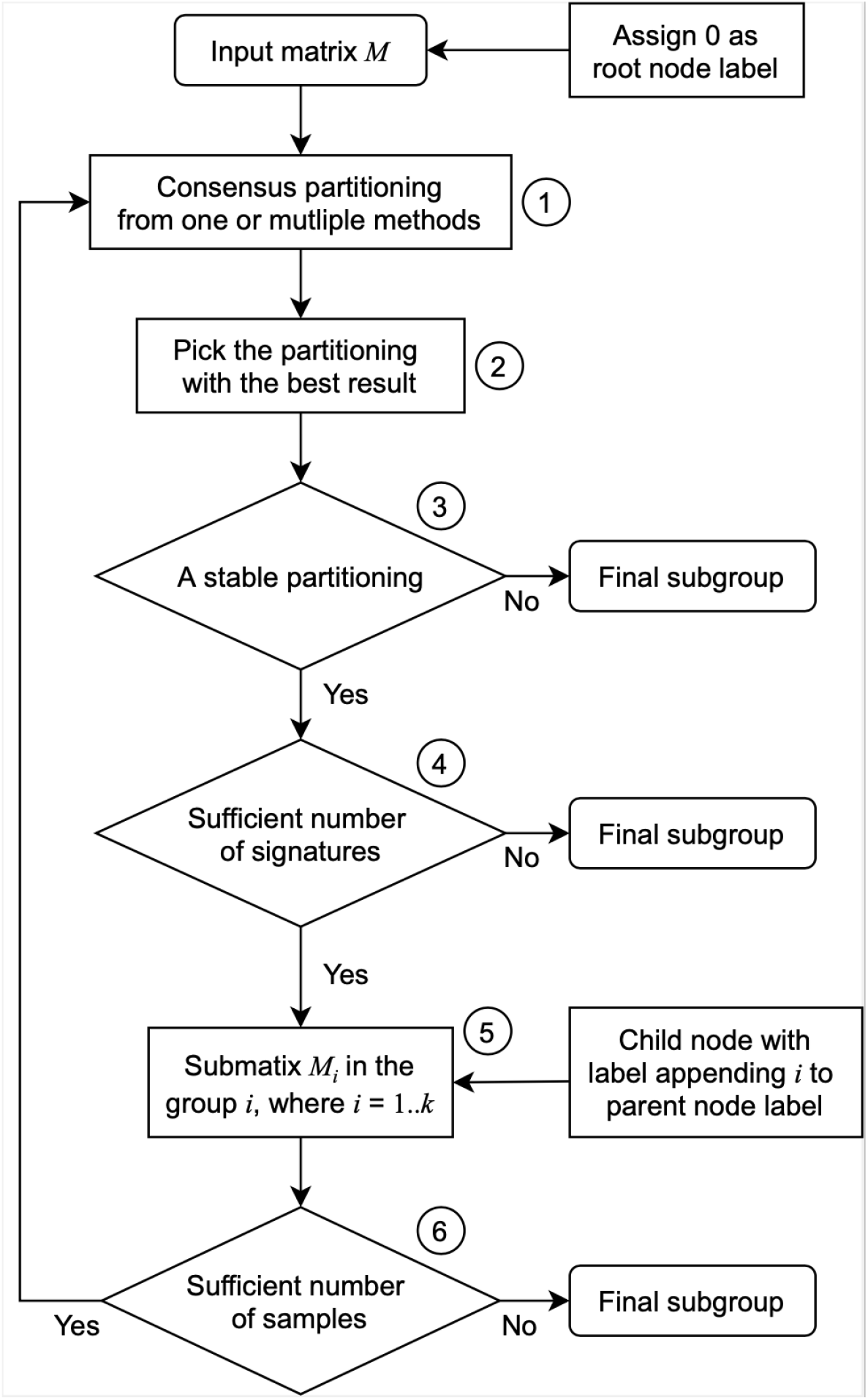
The workflow of hierarchical consensus partitioning. Steps are as follows: 1. Apply consensus partitioning to the data matrix with one combination of top-value method and partitioning method or multiple combination of methods. 2. Pick the partitioning that shows the best result. 3. Test whether the best partitioning is stable. 4. Test whether the best partitioning gives a sufficient number of signatures. 5. If criterions in step 3 and 4 are passed, each subgroup in the partitioning is taken as a child node in HCP. 6. For each submatrix, test whether the number of columns are sufficient. If yes, HCP is applied to the child node recursively.

In HCP, node labels are encoded as a list of digits in a special way. The number of digits corresponds to the depth of the node in the hierarchy and the value of the digits corresponds to the subgroup index in the current node. *E*.*g*., a label of “012” means the node is the second subgroup of the partition that comes from the first subgroup of the complete dataset.

### Automatically selecting the number of top features

In each iteration of CP applied on the submatrix, top features are firstly selected. Generally, secondary subgroups have smaller numbers of features efficient for classification, thus setting the same number of top features for all submatrice would bring additional noises and unstabilize the classification for secondary subgroups. Therefore, a method is needed for automatically selecting a proper number of top features on each step in HCP. This is basically a task of selecting a proper cutoff for filtering the top values. A reasonable way is to select the “elbow” of the top value curve if top values are sorted increasingly. Here we used the method proposed in (Satopaa et al. 2011). It selects the point that has the largest vertical offset to the straight line that connects the points with the minimal top value and the maximal top value. More details can be found in Supplementary File 2.

### Consensus partitioning with down-sampling for large datasets

CP is by-nature a time-consuming analysis. For large datasets with huge numbers of samples, in early steps of the hierarchical procedure, numbers of samples in the subsets could still be large. To improve the efficiency of partitioning on large datasets, we propose a strategy that only applies CP to a small subset of samples that are uniformly picked from the complete submatrix. Later the class labels of unselected samples are predicted by the classification from selected samples. The prediction is based on the signature centroid matrix. For a selected *k*, the signatures that significantly discriminate *k* subgroups are extracted. The signature centroid matrix is a *k*-column matrix where each column is the centroid of confident samples (with silhouette score > 0.5) in the corresponding subgroup, *i*.*e*., the mean across samples. The class prediction is applied as follows: For each unselected sample, we test which signature centroid the current sample is the closest to. For the vector denoted as ***x*** which corresponds to an unselected sample, to predict the class label, the distance calculated by *e*.*g*. Euclidean, cosine or correlation methods to all *k* signature centroids are calculated and denoted as *d*_1_, *d*_2_, …, *d*_*k*_. The class with the smallest distance is assigned to sample:

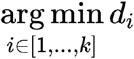

Only using a small subset of samples for classification might completely miss samples in small subgroups, but they can be assigned to the subgroup that they are closest to and they can be identified in later steps of HCP.

In the vignette of the *cola* package, we additionally propose a method that calculates *p*-values for the class label assignment by permuting rows of the signature centroid matrix. It provides confidence for the class label assignment, however, the *p*-value calculation is ignored in the process of HCP because all unselected samples are assigned to the corresponding subgroups regardless of their confidence.

### Compare two classifications

In the Results section, we compared HCP classification to the one from original studies. Here we define similarity measurement for two classifications. For two classifications denoted as *C*_1_ and *C*_2_ with number of subgroups *n*_1_ and *n*_2_ respectively, let *g*_*i*_ and *h*_*j*_ be the *i*th group in *C*_1_ and the *j*th group in *C*_2_. Since different classification methods can hardly generate consistent classifications for all subgroups, especially when *n*_1_ and *n*_2_ are large, we treat *g*_*i*_ to agree with *h*_*j*_ if *g*_*i*_ is fully covered by *h*_*j*_, or vice versa, thus the overlap coefficient is used to measure the similarity of subgroups of two classifications. Assume *A* as the set of samples of *g*_*i*_ and *B* as the set of samples of *h*_*j*_, the overlap coefficient denoted as *a*_*i,j*_ is defined as:

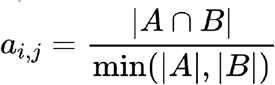

Where |*A*| and |*B*| are the number of samples in the two sets. The agreement of *g*_*i*_ to *C*_2_ is calculated as the maximal similarity to all subgroups in *C*_2_:

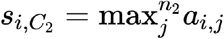

Then the overall similarity of *C*_1_ to *C*_2_ is calculated as the mean of the agreement of each subgroup to *C*_2_ weighted by the subgroup size:

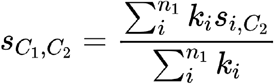

where *k*_*i*_ is the size of *g*_*i*_. We name it *classification agreement* in the paper. The definition of the classification agreement is not exactly symmetric, *i*.*e*. 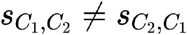, but the two values are very similar.

### Implementation of hierarchical consensus partitioning in *cola*

HCP has been integrated in *cola* from version 2.0.0 under an object-oriented implementation. The main function `hierarchical_partition()` performs the analysis and it returns a *HierarchicalPartition* object. *cola* provides rich visualization utilities on it and we try to implement the application programming interface (API) for the functions the same as those in standard *cola* analysis to make it seamless to switch analysis methods. To name a few: `collect_classes()` draws the hierarchy of the classification; `get_signatures()` calculates and visualizes the rows that are significantly different between subgroups; `dimension_reduction()` performs dimension reduction analysis to visualize how well the subgroups are separated; `top_rows_overlap()` compares the top features on each node; And `functional_enrichment()` automatically applies function enrichment on the signatures if they can be corresponded to genes.

Similar to standard CP analysis in *cola*, there is also a `cola_report()` function that applies on the *HierarchicalPartition* object and automatically performs the complete analysis and generates all the plots in a HTML report. Thus, to perform a hierarchical consensus partitioning analysis, users only need a minimal set of code of using two functions, such as:

~~~
rh = hierarchical_partition(matrix, …)
cola_report(rh, …)
~~~

In Figure 1, step 3, 4, and 6 validate whether HCP should continue on the current node. The validation can also be applied after the classification hierarchy is generated. In most functions, users can control what level in the hierarchy they want by adjusting by the number of samples and number of signatures. Also the dendrogram can be manually merged by `merge_node()` and extended by `split_node()` functions on specific nodes.

### Pre-processing test datasets

The HSMM single-cell RNA-Seq dataset is available in the *HSMMSingleCell* Bioconductor package (Trapnell 2021). Expression values were normalized by log_10_(FPKM + 1) and only the protein-coding genes were used. The peripheral blood mononuclear cells (PBMC) dataset as well as the *Seurat* classification (Satija et al. 2015) were obtained according to the protocol in the tutorial of the *Seurat* package (https://satijalab.org/seurat/articles/pbmc3k_tutorial.html). The central nervous system tumors (CNST) dataset (Capper et al. 2018) was downloaded from the GEO database with accession ID GSE90496. The source and processing of the 66 test datasets can be found in Supplementary File 3 with runnable code.

## Results

### Compare to standard consensus partitioning - a case study

In Figure 2, we demonstrated that CP was difficult to simultaneously identify a large number of subgroups with the HSMM dataset (Trapnell 2021). Here we applied HCP on the same dataset. For CP applied on every node represented as a subset of samples, ATC was set as the top-value method and skmeans was set as the partitioning method.

HCP identified 12 subgroups which were well separated in the *t*-SNE plot (Figure 4A), as a comparison, CP only suggested 6 as the optimized number of subgroups (Figure 2). An overlap of the top features extracted on every node in the subgroup hierarchy showed each subset of samples had its own specific features (Figure 4B). *E*.*g*. on node “02”, 50.1% out of the 1139 top features were unique to all other nodes (Figure 4B). The signature heatmap (Figure 4C) which included the genes with significant differences in at least two subgroups showed the 12 subgroups were well separated and had distinct patterns. The functional enrichment on the signatures also showed they were biologically meaningful, *e*.*g*. genes in row group “km1” were enriched with functions related to cell development and differentiation and genes in row group “km2” were enriched with functions of cell cycle (Figure 4D). HCP involves multiple executions of CP on each node where resampling was randomly applied. Figure 4E demonstrated that randomness had almost no influence on the classification, where HCP was executed 20 times and the classification showed high stability with 97.9% concordance (measured as the average percent of samples having the same classification).

**Figure 4.**
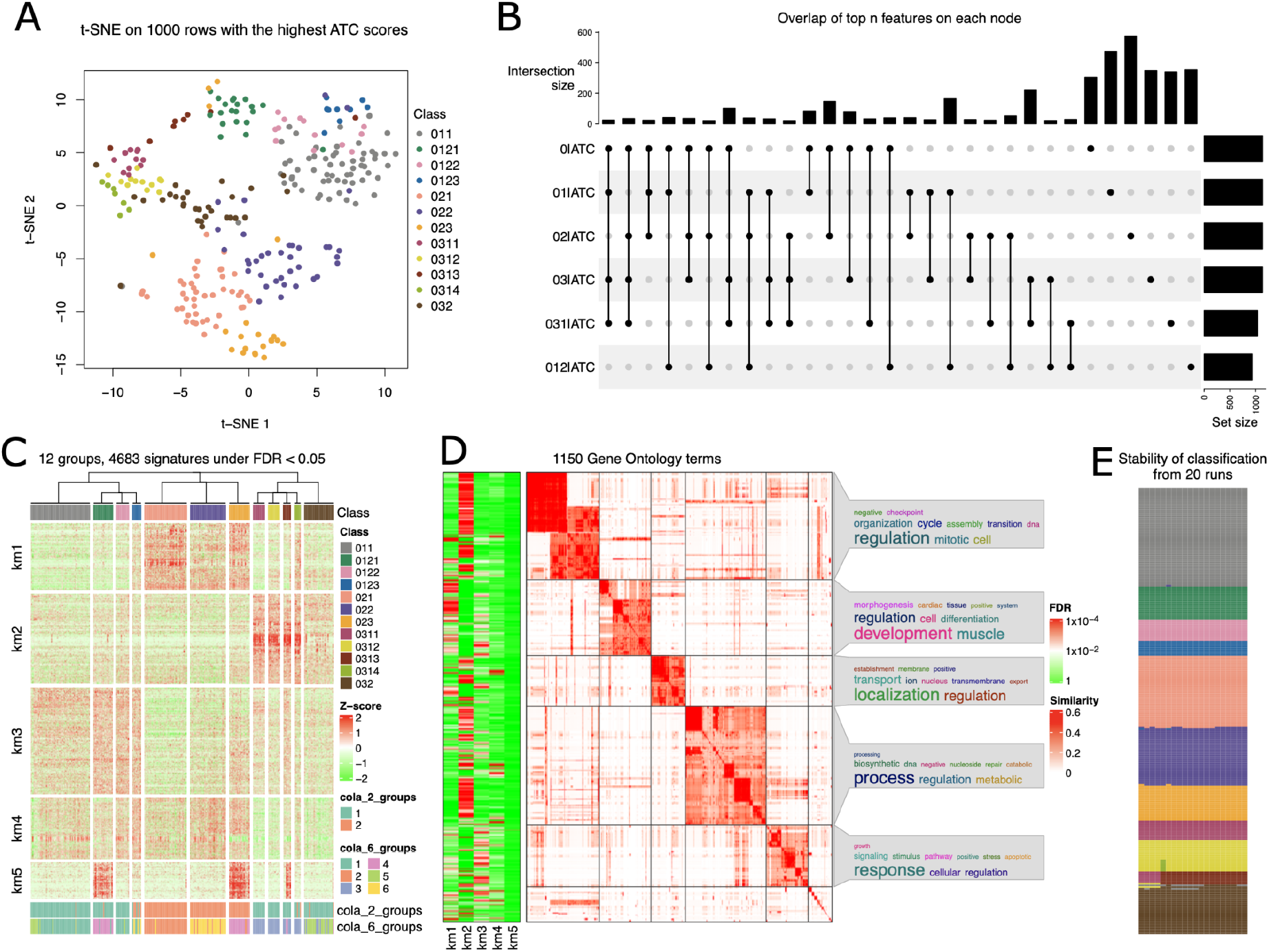
Application of HCP on the HSMM dataset. A) *t*-SNE plot of the top 1000 genes with the highest ATC scores. B) The UpSet plot showing the overlap of top features selected on each node of HCP. Only combinations with size larger than 20 are shown in the plot. C) Heatmap of the signature genes under HCP classification. The dendrogram on top of heatmap corresponds to the hierarchy of the HCP classification. D) Gene Ontology (GO) enrichment of the signature genes. The left heatmap visualizes the FDRs from the enrichment analysis and the right heatmap visualizes the similarity of GO terms where the GO terms are clustered by their similarities. The analysis was performed with the *simplifyEnrichment* package. E) Stability of the HCP classification from 20 repetitive runs.

### Applications on a large scRNAseq dataset

We applied HCP on the PBMC scRNASeq dataset that contains 2638 cells (Satija et al. 2015). In HCP, when the subset on a node had more than 500 cells, down sampling was turned on. ATC was used as the top-value method and skmeans was used as the partitioning method. Branching of HCP stopped when the number of signatures was less than 200. We compared HCP classification to CP classification as well as the original classification analyzed with the *Seurat* package (Satija et al. 2015).

HCP identified 7 subgroups (Figure 5A) and CP only identified 4 subgroups (Figure 5C) as the optimized result, where group “2” in CP was additionally classified into three subgroups in HCP which are “0211”, “”0212” and “022” and group labeled “3” in CP was additionally classified into two subgroups in HCP labeled “041” and “042”. According to the points distribution in Figure 5C, there were indeed four major subgroups that can be observed, but HCP can reveal more secondary subgroups.

**Figure 5.**
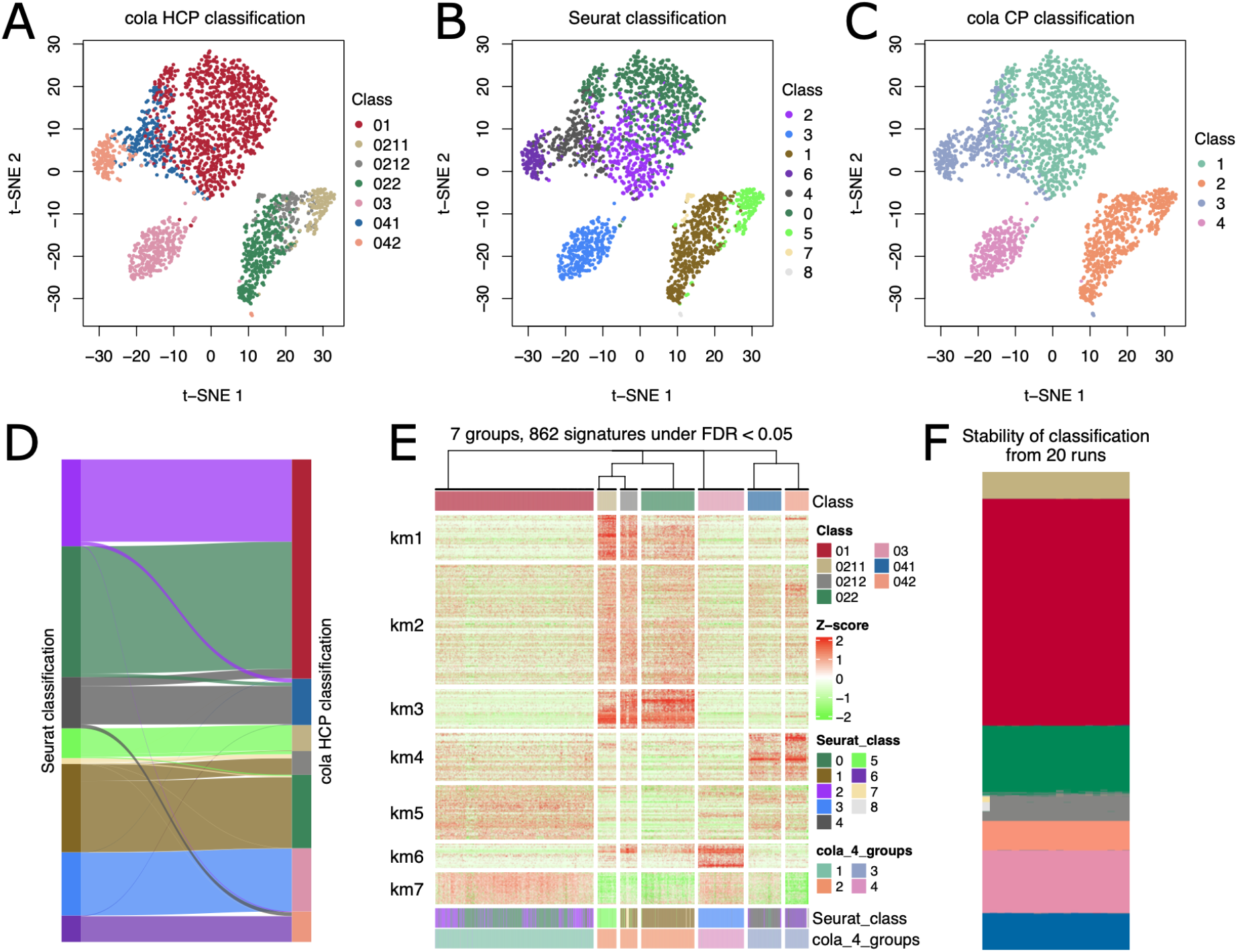
Application of HCP on PBMC scRNAseq dataset. A-C) *t*-SNE plot of the top 1000 genes with the highest ATC scores. The coloring is based on HCP classification, *Seurat* classification and CP classification. D) Correspondence between HCP and *Seurat* classifications. E) Signature genes from HCP classification. F) Stability of the HCP classifications form 20 repetitive runs.

When Comparing HCP to *Seurat* classification, most samples had similar classifications except group “01” in HCP was split into two groups under *Seurat* (“0”/”2”) and some disagreement between HCP “0212”/”022” and *Seurat* “1”/“7” (Figure 5A, 5B, 5D). The overall classification agreement between the two classifications is 0.947. In Supplementary File 4, we demonstrated that node “01” can be further split into two sub-nodes (“011”/”012”) but with less stability, where classification of “011”/”012” was similar to *Seurat* “0”/“2”, and this classification separated the two groups more than *Seurat* classification. We also compared the HCP “0212”/”022” and Seurat “1”/”7” in Supplementary File 4 and we found the two classifications on this subset of samples generated different sets of signature genes and signature genes from HCP classification had more significant biological functions.

For this dataset, random sampling was additionally applied when the number of nodes was larger than 500, which brought a second layer of randomness. Nevertheless, Figure 5F illustrates the classification is stable, which implies that if there exists a clear classification, classification by down-sampling won’t bring significant noise from randomization (concordance of the classifications from 20 repetitive runs is 99.1%).

### Applications on a methylation dataset with a large number of subgroups

We applied HCP on a DNA methylation array dataset of central nervous system tumors (CNST) with 2803 samples (Capper et al. 2018). The dataset contains 14 different tumor types (including controls) which were additionally classified into 91 subtypes based on methylation profiles (The correspondence between tumor types and methylation classes is illustrated in Supplementary File 5). The aim of this analysis is to see whether HCP can recover such a great number of subgroups. In the analysis, we only considered CpG probes located in CpG islands. One reason is for reducing the dataset and the other reason is we have demonstrated it might be more proper to analyze CpGs in different CpG features since they might generate different classifications and correspond to different biological meanings (Gu et al. 2020). The final matrix for HCP analysis contains 17976 probes. In the analysis, the configurations are as follows: for CP executed on each node, two top-value methods (SD and ATC) and two partitioning methods (kmeans and skmeans) were tested because SD/kmeans are popular choice in current studies and we demonstrated ATC/skmeans were better for identifying subgroups, thus we used the 4 combinations of methods for CP on nodes and HCP automatically picked the one with the best result on each node. Rows were not scaled because methylation data was all in the same scale, *i*.*e*., 0∼1. In each submatrix, a filtering step was applied ahead of clustering which only took the top 30000 probes with the highest SD values. Later top 1000 features extracted by SD/ATC from 30000 probes were only used for partitioning. This pre-filtering is for reducing the running time of ATC score calculation. To get rid of ATC extracting rows showing high correlation but with small absolute differences, the rows difference of methylation between subgroups for signature probes was set to > 0.25. And finally, the minimal number of signature probes was set to > 1000. When the number of samples in a submatrix is larger than 500, down sampling was enabled.

HCP identified in total 92 subgroups. In the original study, samples were classified into 14 different tumor types. Figure 6A demonstrates the agreement between tumor types and HCP classifications. It shows in most cases, most of the samples in the same HCP subgroup always belong to a single tumor type, except in a few cases, samples in the same HCP subgroup belong to multiple tumor types. The overall classification agreement between HCP classification and tumor types is 0.903. Only 15/92 (16.3%) HCP subgroups (covering 15.2% of all samples) are less compatible with tumor types with overlap coefficient < 0.75 (Figure 6C).

**Figure 6.**
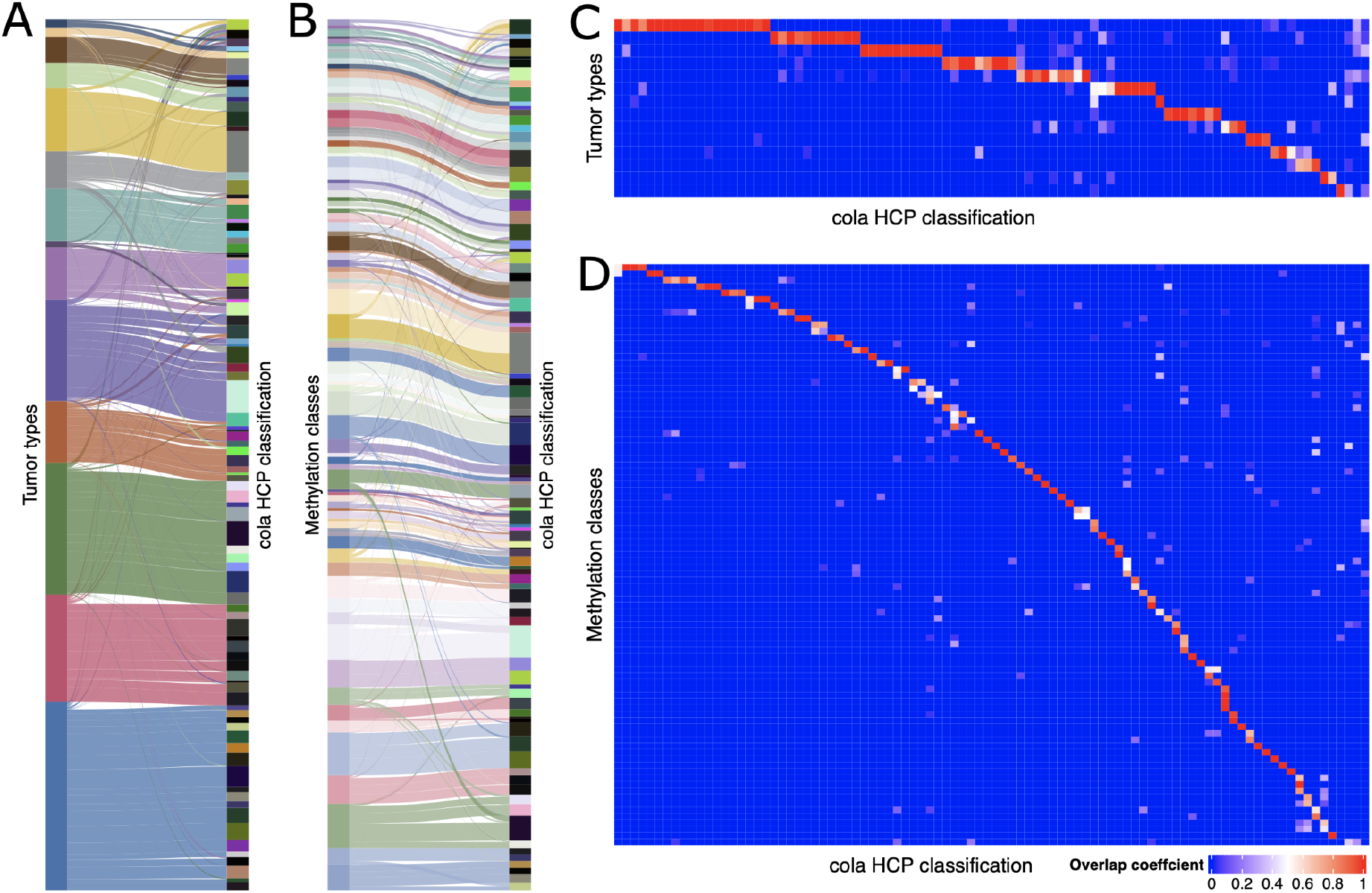
Compare classifications on the CNST dataset. A) Compare classification between tumor types and *cola* HCP classification. B) Compare classifications between methylation classes and *cola* HCP classifications. C). Overlap coefficient between tumor types and *cola* HCP classification. D) overlap coefficient between methylation class and *cola* HCP classifications.

The 14 tumor types were additionally classified into 91 subtypes based on methylation profile in the original study (Supplementary File 5). Figure 6B illustrates the correspondence between HCP classification and methylation classifications. The overall classification agreement is 0.868 and only 17/92 (18.5%) HCP subgroups (covering 16.5% of all samples) were less compatible with methylation classes with overlap coefficient < 0.75 (Figure 6D).

We can also find 29 HCP subgroups had a one-to-one mapping to methylation classes, 11 methylation classes that covered multiple HCP subgroups, and 6 HCP subgroups covered multiple methylation classes, with overlap coefficient > 0.75 (Figure 6D).

## Discussion

Nowadays, as sample sizes of cohort studies increase fastly, it provides possibilities for detecting more subtle subgroups that have more specific patterns in the data. In tumor studies, subtypes are frequently studied (Ceccarelli et al. 2016) (Cancer Genome Atlas Research Network et al. 2016) (Liu et al. 2018). However, due to the complex mechanisms of tumor generations, *e*.*g*., tumor microenvironment, inter/intra-cell interactions, spatial as well as temporal patterns, it makes tumors with more specific molecular profiles, thus, it is important to identify these subtypes with highly specific profiles, which benefits a more precise diagnosis for tumors (Ogino et al. 2012; Rich 2016). In single cell technologies, now it allows one to simultaneously measure a great number of cells, which gives a chance for detecting the hierarchy of various cell types (Liberzon et al. 2015). Due to the nature of the biological system, it is believed they have a hierarchical structure. For consensus partitioning methods, although they have been successfully applied to reveal tumor subtypes, however, they are still weak for identifying more subtypes which show different levels of differences. A classification with hierarchical methods can solve this problem (Silla and Freitas 2011). In this work, we proposed a new method which applies consensus partitioning in a hierarchy procedure. It integrates the advantages of having the stability of classification as well as revealing subgroups with multi-level and subtle differences. We applied HCP on real-world dataset and it shows HCP can efficiently reveal more subtypes and have more meaningful classifications compared to current ones. In Supplementary File 3, we additionally demonstrated the use of HCP with 66 real-world datasets.

HCP has a weakness that misclassifications in early steps in the hierarchical process will accumulate and affect the downstream classifications. One scenario for this misclassification is when secondary subgroups have less consistent patterns, samples inside are partially assigned to different subgroups, and once they are separated, they cannot be merged back in later steps of HCP. One solution is to increase maximal *k* tried on each node and the other solution is to try various partitioning methods simultaneously on the node where the partitioning with misclassification always has less stability and can be filtered out. However, this can only be a problem when the patterns are not clear.

Consensus partitioning is a multiple-step analysis where selection of parameters on each step might affect the final partitioning result. HCP involves a list of executions of CP on nodes of the subgroup hierarchy, thus, CP should be applied in a way that parameters are optimally selected to ensure that subgroups are successfully separated on each node and hierarchy is sufficiently extended. In HCP, several strategies were implemented. The number of top features for each submatrix was automatically selected, in order to exclude those features that might bring additional noise for classification. Also the silhouette score instead of 1-PAC score was used to validate the stability of classification because silhouette score provides a stricter way to measure the stability and it prefers to select a smaller but more stable *k*, and more subgroups will be found in the later steps of the hierarchical procedures.

Highly heterogeneous data might result in deep iterations in the hierarchical process and generate large numbers of subgroups. As demonstrated in the analysis of the TCGA GBM microarray dataset (Supplementary File 3 and 6), standard CP generated 4 subgroups as the best result while HCP generated 16 subgroups where the mean subgroup size was only 11. Although on one hand this can be an example of standard CP not being able to detect a large number of subgroups, on the other hand it shows HCP might over-classify samples and lose generality of the classification. Indeed, the separation of the 16 subgroups by HCP was biologically reasonable where large numbers of signature genes on all nodes (> 600 under FDR < 0.05) supported the classification on the corresponding nodes, but too specific classification would increase the difficulty to Interpret the results and to extend to different studies based on the same biological objects. To balance the generality and specificity, it depends on what level of the heterogeneity users expect. A simple solution is to set the minimal number of samples on nodes. The other solution is to filter the hierarchy by the number of signatures on each node, which provides a view of the biological importance of the classification and generally the number of signatures decreases when subgroups have smaller separation. To facilitate users to adjust the level of heterogeneity of classifications, HCP in *cola* provides functionalities to extract and analyze the classification at a specific level of subgroup hierarchy.

When the numbers of samples were large, we proposed to apply CP to a subset of randomly sampled samples. Later the class labels of the unselected samples were predicted. If there are clear separations of major subgroups, downsampling CP performs well enough that it retains the classification of major groups, while if classification is not stable for the complete set of samples, classification on the randomly selected subset tends to be unstable. Thus, down sampling is a good strategy for data with clear structures to reduce running time. Downsampling might miss samples in small subgroups, however, they can be attached to major subgroups and be revealed in later steps of HCP, thus downsampling CP is a good companion of HCP.

## Conclusion

In this work, we proposed a new method that improves consensus partitioning via a hierarchical procedure. It can reveal more subgroups with various levels of differences which are weak to detect by standard consensus partitioning methods. We demonstrated the usage of hierarchical consensus partitioning with real-world examples. We also demonstrated its ability to identify great numbers of subgroups with high agreement to the original studies. The method has been implemented in the *cola* package. The functionality of hierarchical consensus partitioning was designed with both ease and comprehensiveness.

We believe it will be a convenient and powerful tool for diving deeper into users’ data to reveal more meaningful results.

## Data Availability

Hierarchical consensus partitioning has been implemented in the *cola* package from version 2.0.0 (https://bioconductor.org/packages/cola/). The HTML reports for 66 test datasets are publicly available at https://cola-rh.github.io/. The scripts to perform the complete analysis are available at https://cola-rh.github.io/manuscript. The Supplementary Files are also available at https://cola-rh.github.io/supplementary/.

## Acknowledgement

This work was supported by the NCT Molecular Precision Oncology Program.

