## Supplementary File 1 for "Improve consensus partitioning via a hierarchical procedure"

Apply hierarchical consensus partitioning on the example dataset in Figure 1


### Apply hierarchical consensus partitioning on the example dataset in Figure 1

###### Zuguang Gu

#### 2021-07-29

In Figure 1 of the manuscript, we constructed a random matrix that contained groups with large difference as well as groups with small difference and we demonstrated that standard consensus partitioning procedures cannot separate all the four groups simultaneously. In this supplementary, we demonstrate that hierarchical consensus partitioning is able to identify all groups.

First we generate the random matrix the same as in Figure 1.

```
library(ComplexHeatmap)

set.seed(54)
mean_diff1 = rnorm(100)

m1 = do.call(rbind, lapply(1:100, function(i) {
    c(rnorm(50, mean = mean_diff1[i]), rnorm(50, mean = -mean_diff1[i]))
}))

mean_diff2 = rnorm(100)/2
mean_diff2[order(abs(mean_diff1))] = mean_diff2[order(abs(mean_diff2), decreasing = TRUE)]

m2 = do.call(rbind, lapply(1:100, function(i) {
    c(rnorm(10, mean = mean_diff2[i]), rnorm(10, mean = -mean_diff2[i]))
}))

m = cbind(m1, m2)

group = rep(c("A1", "A2", "B1", "B2"), times = c(50, 50, 10, 10))
group_col = structure(c("#E41A1C", "#377EB8", "#4DAF4A", "#984EA3"), names = c("A1", "A2", "B1", "B2"))

Heatmap(m, name = "mat", 
    top_annotation = HeatmapAnnotation(Group = group, col = list(Group = group_col)),
    show_row_dend = FALSE, column_title = "Heatmap of the random dataset",
    row_dend_reorder = mean_diff1, column_dend_reorder = as.numeric(factor(group))
)
```

Figure S1.1. Heatmap of the random matrix used in Figure 1 in the manuscript.

The standard consensus partition procedures cannot identify all four groups. It can only identify three groups as the best results:

```
res = consensus_partition(m, top_value_method = "SD", partition_method = "kmeans",
    top_n = 50, anno = group, anno_col = group_col, scale_rows = FALSE)
collect_plots(res)
```

Figure S1.2. Various plots for interpreting consensus partitioning results.

Then we apply hierarchical consensus partitioning with the function `hierarchical_partition()` on the matrix:

```
library(cola)
rh = hierarchical_partition(m, 
    top_value_method = "SD", partition_method = "kmeans", 
    anno = group, anno_col = group_col,
    top_n = 50, scale_rows = FALSE)
```

We can print the `rh` object:

```
rh
```

```
## A 'HierarchicalPartition' object with 'SD:kmeans' method.
##   On a matrix with 100 rows and 120 columns.
##   Performed in total 900 partitions.
##   There are 4 groups under the following parameters:
##     - min_samples: 6
##     - mean_silhouette_cutoff: 0.9
##     - min_n_signatures: 4 (signatures are selected based on:)
##       - fdr_cutoff: 0.05
##       - group_diff: 0
## 
## Hierarchy of the partition:
##   0, 120 cols
##   |-- 01, 50 cols (a)
##   |-- 02, 50 cols (a)
##   `-- 03, 20 cols, 27 signatures
##       |-- 031, 10 cols (b)
##       `-- 032, 10 cols (b)
## Stop reason:
##   a) Mean silhouette score was too small
##   b) Subgroup had too few columns.
## 
## Following methods can be applied to this 'HierarchicalPartition' object:
##  [1] "all_leaves"            "all_nodes"             "cola_report"          
##  [4] "collect_classes"       "colnames"              "compare_signatures"   
##  [7] "dimension_reduction"   "functional_enrichment" "get_anno"             
## [10] "get_anno_col"          "get_children_nodes"    "get_classes"          
## [13] "get_matrix"            "get_signatures"        "is_leaf_node"         
## [16] "max_depth"             "merge_node"            "ncol"                 
## [19] "node_info"             "node_level"            "nrow"                 
## [22] "rownames"              "show"                  "split_node"           
## [25] "suggest_best_k"        "test_to_known_factors" "top_rows_heatmap"     
## [28] "top_rows_overlap"     
## 
## You can get result for a single node by e.g. object["01"]
```

The function `collect_classes()` draws the subgroup hierarchy:

```
collect_classes(rh)
```

Figure S1.3. Hierarchiy of cola HCP.

As we can see, all the four groups are separated.

#### Session info

```
sessionInfo()
```

```
## R version 4.1.0 (2021-05-18)
## Platform: x86_64-apple-darwin17.0 (64-bit)
## Running under: macOS Big Sur 10.16
## 
## Matrix products: default
## BLAS:   /Library/Frameworks/R.framework/Versions/4.1/Resources/lib/libRblas.dylib
## LAPACK: /Library/Frameworks/R.framework/Versions/4.1/Resources/lib/libRlapack.dylib
## 
## locale:
## [1] C/UTF-8/C/C/C/C
## 
## attached base packages:
##  [1] parallel  stats4    grid      stats     graphics  grDevices utils    
##  [8] datasets  methods   base     
## 
## other attached packages:
##  [1] simplifyEnrichment_1.3.1 org.Hs.eg.db_3.13.0      AnnotationDbi_1.54.1    
##  [4] IRanges_2.26.0           S4Vectors_0.30.0         Biobase_2.52.0          
##  [7] BiocGenerics_0.38.0      eulerr_6.1.0             cowplot_1.1.1           
## [10] genefilter_1.74.0        GetoptLong_1.0.5         ComplexHeatmap_2.9.3    
## [13] circlize_0.4.13          cola_1.9.4               knitr_1.33              
## [16] rmarkdown_2.9            BiocManager_1.30.16      colorout_1.2-2          
## 
## loaded via a namespace (and not attached):
##   [1] shadowtext_0.0.8       fastmatch_1.1-0        plyr_1.8.6            
##   [4] igraph_1.2.6           lazyeval_0.2.2         proxyC_0.2.0          
##   [7] polylabelr_0.2.0       splines_4.1.0          Polychrome_1.2.6      
##  [10] BiocParallel_1.26.0    GenomeInfoDb_1.28.0    ggplot2_3.3.5         
##  [13] digest_0.6.27          foreach_1.5.1          htmltools_0.5.1.1     
##  [16] GOSemSim_2.18.0        viridis_0.6.1          magick_2.7.2          
##  [19] GO.db_3.13.0           fansi_0.5.0            magrittr_2.0.1        
##  [22] memoise_2.0.0          tm_0.7-8               cluster_2.1.2         
##  [25] doParallel_1.0.16      Biostrings_2.60.1      annotate_1.70.0       
##  [28] graphlayouts_0.7.1     RcppParallel_5.1.4     matrixStats_0.59.0    
##  [31] enrichplot_1.12.1      colorspace_2.0-2       blob_1.2.1            
##  [34] ggrepel_0.9.1          xfun_0.24              dplyr_1.0.7           
##  [37] crayon_1.4.1           RCurl_1.98-1.3         microbenchmark_1.4-7  
##  [40] jsonlite_1.7.2         scatterpie_0.1.6       impute_1.66.0         
##  [43] ape_5.5                brew_1.0-6             survival_3.2-11       
##  [46] iterators_1.0.13       glue_1.4.2             polyclip_1.10-0       
##  [49] gtable_0.3.0           zlibbioc_1.38.0        XVector_0.32.0        
##  [52] shape_1.4.6            scales_1.1.1           DOSE_3.18.1           
##  [55] data.tree_1.0.0        bezier_1.1.2           DBI_1.1.1             
##  [58] Rcpp_1.0.6             gridtext_0.1.4         viridisLite_0.4.0     
##  [61] xtable_1.8-4           clue_0.3-59            tidytree_0.3.4        
##  [64] bit_4.0.4              mclust_5.4.7           httr_1.4.2            
##  [67] fgsea_1.18.0           RColorBrewer_1.1-2     ellipsis_0.3.2        
##  [70] pkgconfig_2.0.3        XML_3.99-0.6           farver_2.1.0          
##  [73] sass_0.4.0             utf8_1.2.1             tidyselect_1.1.1      
##  [76] rlang_0.4.11           reshape2_1.4.4         munsell_0.5.0         
##  [79] tools_4.1.0            cachem_1.0.5           downloader_0.4        
##  [82] generics_0.1.0         RSQLite_2.2.7          evaluate_0.14         
##  [85] stringr_1.4.0          fastmap_1.1.0          yaml_2.2.1            
##  [88] ggtree_3.0.2           bit64_4.0.5            tidygraph_1.2.0       
##  [91] purrr_0.3.4            dendextend_1.15.1      KEGGREST_1.32.0       
##  [94] ggraph_2.0.5           nlme_3.1-152           slam_0.1-48           
##  [97] aplot_0.0.6            DO.db_2.9              xml2_1.3.2            
## [100] compiler_4.1.0         png_0.1-7              treeio_1.16.1         
## [103] tibble_3.1.2           tweenr_1.0.2           bslib_0.2.5.1         
## [106] stringi_1.6.2          highr_0.9              lattice_0.20-44       
## [109] Matrix_1.3-4           markdown_1.1           vctrs_0.3.8           
## [112] pillar_1.6.1           lifecycle_1.0.0        jquerylib_0.1.4       
## [115] GlobalOptions_0.1.2    data.table_1.14.0      bitops_1.0-7          
## [118] irlba_2.3.3            patchwork_1.1.1        qvalue_2.24.0         
## [121] R6_2.5.0               gridExtra_2.3          codetools_0.2-18      
## [124] MASS_7.3-54            assertthat_0.2.1       rjson_0.2.20          
## [127] GenomeInfoDbData_1.2.6 clusterProfiler_4.0.0  tidyr_1.1.3           
## [130] rvcheck_0.1.8          skmeans_0.2-13         Cairo_1.5-12.2        
## [133] ggforce_0.3.3          scatterplot3d_0.3-41   NLP_0.2-1
```
