## Supplementary File 2 for "Improve consensus partitioning via a hierarchical procedure"

Automatically select the ‘knee/elbow’ of a curve


### Automatically select the ‘knee/elbow’ of a curve

###### Zuguang Gu

#### 2021-07-29

We demonstrate the method that automatically selects the “knee/elbow” of a curve proposed in the study Satoppa et al., 2011.

We first get a vector of scores which are the row means of the matrix from the Golub dataset.

```
library(cola)
library(golubEsets)
data(Golub_Merge)
m = exprs(Golub_Merge)

m[m <= 1] = NA
m = log10(m)
m = adjust_matrix(m)

library(matrixStats)
s = rowSds(m)
```

Values in `s` are sorted increasingly and they are visualized as a scatterplot. Now the task is to identify the “elbow” of this curve.

```
s = sort(s)
plot(s)
```

Figure S2.1. Sorted row means of Golub dataset.

Satoppa et al., 2011 proposed a method that selects the “elbow point” as the one having the largest distance to the straight line (denoted as \(L\)) that connects the two boundary points with the minimal and maximal values on *y*-axis. The method is demonstrated by the following code. Note values on both *x*-axis and *y*-axis are scaled into `[0, 1]`.

```
y = s
x = seq_along(y)

x = x/max(x)
y = y/max(y)

n = length(x)
a = (y[n] - y[1])/(x[n] - x[1])
b = y[1] - a * x[1]
d = a * x + b - y
x1 = x[which.min(d)]
x2 = x[which.max(d)]
theta = atan(a)

plot(x, y, xlab = "index", ylab = "value", asp = 1)
abline(a = b, b = a)

breaks = seq(1000, 3800, length = 8)/n
y0 = quantile(y, breaks)
y1 = a*breaks + b
segments(breaks, y0, breaks, y1, col = "red")

a2 = tan(theta + pi/2)
b2 = y0- a2*breaks
x2 = (b2 - b)/(a - a2)
y2 = a*x2 + b
segments(breaks, y0, x2, y2, col = "blue")
```

Figure S2.2. Demonstration of the method to look for elbow/knee of the curve.

Instead of selecting by the distance to \(L\) (the lengths of blue lines), we can simply select according to the vertical distance to \(L\) (the lengths of red lines). The two selection rules are actually identical.

We implement this method with the function `knee_finder2()` in **cola**. The function generates two plots: the original curve and a curve of the vertical distance to \(L\).

```
knee_finder2(s, plot = TRUE)
```

Figure S2.3. Output plots from knee\_finder2().

```
## [1]   NA 3790
```

`knee_finder()` returns a vector of two. The second value corresponds to the elbow of the curve and the first value corresponds to the knee of the curve, as shown in the following plot.

```
s = rnorm(1000)
knee_finder2(s, plot = TRUE)
```

Figure S2.4. When both knee and elbow exist in the curve.

```
## [1] 107 888
```

#### Session info

```
sessionInfo()
```

```
## R version 4.1.0 (2021-05-18)
## Platform: x86_64-apple-darwin17.0 (64-bit)
## Running under: macOS Big Sur 10.16
## 
## Matrix products: default
## BLAS:   /Library/Frameworks/R.framework/Versions/4.1/Resources/lib/libRblas.dylib
## LAPACK: /Library/Frameworks/R.framework/Versions/4.1/Resources/lib/libRlapack.dylib
## 
## locale:
## [1] C/UTF-8/C/C/C/C
## 
## attached base packages:
##  [1] parallel  stats4    grid      stats     graphics  grDevices utils    
##  [8] datasets  methods   base     
## 
## other attached packages:
##  [1] matrixStats_0.59.0       golubEsets_1.34.0        simplifyEnrichment_1.3.1
##  [4] org.Hs.eg.db_3.13.0      AnnotationDbi_1.54.1     IRanges_2.26.0          
##  [7] S4Vectors_0.30.0         Biobase_2.52.0           BiocGenerics_0.38.0     
## [10] eulerr_6.1.0             cowplot_1.1.1            genefilter_1.74.0       
## [13] GetoptLong_1.0.5         ComplexHeatmap_2.9.3     circlize_0.4.13         
## [16] cola_1.9.4               knitr_1.33               rmarkdown_2.9           
## [19] BiocManager_1.30.16      colorout_1.2-2          
## 
## loaded via a namespace (and not attached):
##   [1] shadowtext_0.0.8       fastmatch_1.1-0        plyr_1.8.6            
##   [4] igraph_1.2.6           lazyeval_0.2.2         proxyC_0.2.0          
##   [7] polylabelr_0.2.0       splines_4.1.0          Polychrome_1.2.6      
##  [10] BiocParallel_1.26.0    GenomeInfoDb_1.28.0    ggplot2_3.3.5         
##  [13] digest_0.6.27          foreach_1.5.1          htmltools_0.5.1.1     
##  [16] GOSemSim_2.18.0        viridis_0.6.1          magick_2.7.2          
##  [19] GO.db_3.13.0           fansi_0.5.0            magrittr_2.0.1        
##  [22] memoise_2.0.0          tm_0.7-8               cluster_2.1.2         
##  [25] doParallel_1.0.16      Biostrings_2.60.1      annotate_1.70.0       
##  [28] graphlayouts_0.7.1     RcppParallel_5.1.4     enrichplot_1.12.1     
##  [31] colorspace_2.0-2       blob_1.2.1             ggrepel_0.9.1         
##  [34] xfun_0.24              dplyr_1.0.7            crayon_1.4.1          
##  [37] RCurl_1.98-1.3         microbenchmark_1.4-7   jsonlite_1.7.2        
##  [40] scatterpie_0.1.6       impute_1.66.0          ape_5.5               
##  [43] brew_1.0-6             survival_3.2-11        iterators_1.0.13      
##  [46] glue_1.4.2             polyclip_1.10-0        gtable_0.3.0          
##  [49] zlibbioc_1.38.0        XVector_0.32.0         shape_1.4.6           
##  [52] scales_1.1.1           DOSE_3.18.1            data.tree_1.0.0       
##  [55] bezier_1.1.2           DBI_1.1.1              Rcpp_1.0.6            
##  [58] gridtext_0.1.4         viridisLite_0.4.0      xtable_1.8-4          
##  [61] clue_0.3-59            tidytree_0.3.4         bit_4.0.4             
##  [64] mclust_5.4.7           httr_1.4.2             fgsea_1.18.0          
##  [67] RColorBrewer_1.1-2     ellipsis_0.3.2         pkgconfig_2.0.3       
##  [70] XML_3.99-0.6           farver_2.1.0           sass_0.4.0            
##  [73] utf8_1.2.1             tidyselect_1.1.1       rlang_0.4.11          
##  [76] reshape2_1.4.4         munsell_0.5.0          tools_4.1.0           
##  [79] cachem_1.0.5           downloader_0.4         generics_0.1.0        
##  [82] RSQLite_2.2.7          evaluate_0.14          stringr_1.4.0         
##  [85] fastmap_1.1.0          yaml_2.2.1             ggtree_3.0.2          
##  [88] bit64_4.0.5            tidygraph_1.2.0        purrr_0.3.4           
##  [91] dendextend_1.15.1      KEGGREST_1.32.0        ggraph_2.0.5          
##  [94] nlme_3.1-152           slam_0.1-48            aplot_0.0.6           
##  [97] DO.db_2.9              xml2_1.3.2             compiler_4.1.0        
## [100] png_0.1-7              treeio_1.16.1          tibble_3.1.2          
## [103] tweenr_1.0.2           bslib_0.2.5.1          stringi_1.6.2         
## [106] highr_0.9              lattice_0.20-44        Matrix_1.3-4          
## [109] markdown_1.1           vctrs_0.3.8            pillar_1.6.1          
## [112] lifecycle_1.0.0        jquerylib_0.1.4        GlobalOptions_0.1.2   
## [115] data.table_1.14.0      bitops_1.0-7           irlba_2.3.3           
## [118] patchwork_1.1.1        qvalue_2.24.0          R6_2.5.0              
## [121] gridExtra_2.3          codetools_0.2-18       MASS_7.3-54           
## [124] assertthat_0.2.1       rjson_0.2.20           GenomeInfoDbData_1.2.6
## [127] clusterProfiler_4.0.0  tidyr_1.1.3            rvcheck_0.1.8         
## [130] skmeans_0.2-13         Cairo_1.5-12.2         ggforce_0.3.3         
## [133] scatterplot3d_0.3-41   NLP_0.2-1
```
