## Supplementary File 3 for "Improve consensus partitioning via a hierarchical procedure"

Test datasets


### Test datasets

###### Zuguang Gu

#### 2021-07-28

| Dataset | Number of columns | Data type | HTML Report | Data source |
| --- | --- | --- | --- | --- |
| HSMM\_single\_cell | 271 | scRNASeq | https://cola-rh.github.io/HSMM\_single\_cell/ | https://bioconductor.org/packages/HSMMSingleCell/ |
| PBMC | 2638 | scRNASeq | https://cola-rh.github.io/PBMC/ | https://satijalab.org/seurat/articles/pbmc3k\_tutorial.html |
| GSE90496 | 2801 | 450K methylation array | https://cola-rh.github.io/GSE90496/ | https://www.ncbi.nlm.nih.gov/geo/query/acc.cgi?acc=GSE90496 |
| Golub\_leukemia | 72 | Microarray, gene expression | https://cola-rh.github.io/Golub\_leukemia/ | https://bioconductor.org/packages/golubEsets/ |
| Ritz\_ALL | 128 | Microarray, gene expression | https://cola-rh.github.io/Ritz\_ALL/ | https://bioconductor.org/packages/ALL/ |
| TCGA\_GBM\_microarray | 173 | Microarray, gene expression | https://cola-rh.github.io/TCGA\_GBM\_microarray/ | https://gdc.cancer.gov/about-data/publications/gbm\_exp |
| BaronPancreas\_mouse | 1886 | scRNASeq | https://cola-rh.github.io/BaronPancreas\_mouse/ | https://bioconductor.org/packages/scRNAseq/ |
| BuettnerESC | 288 | scRNASeq | https://cola-rh.github.io/BuettnerESC/ | https://bioconductor.org/packages/scRNAseq/ |
| DarmanisBrain | 466 | scRNASeq | https://cola-rh.github.io/DarmanisBrain/ | https://bioconductor.org/packages/scRNAseq/ |
| GrunHSC | 1915 | scRNASeq | https://cola-rh.github.io/GrunHSC/ | https://bioconductor.org/packages/scRNAseq/ |
| GrunPancreas | 1728 | scRNASeq | https://cola-rh.github.io/GrunPancreas/ | https://bioconductor.org/packages/scRNAseq/ |
| KolodziejczykESC | 704 | scRNASeq | https://cola-rh.github.io/KolodziejczykESC/ | https://bioconductor.org/packages/scRNAseq/ |
| LaMannoBrain\_human\_es | 1715 | scRNASeq | https://cola-rh.github.io/LaMannoBrain\_human\_es/ | https://bioconductor.org/packages/scRNAseq/ |
| LaMannoBrain\_human\_embryo | 1977 | scRNASeq | https://cola-rh.github.io/LaMannoBrain\_human\_embryo/ | https://bioconductor.org/packages/scRNAseq/ |
| LaMannoBrain\_human\_ips | 337 | scRNASeq | https://cola-rh.github.io/LaMannoBrain\_human\_ips/ | https://bioconductor.org/packages/scRNAseq/ |
| LaMannoBrain\_mouse\_adult | 243 | scRNASeq | https://cola-rh.github.io/LaMannoBrain\_mouse\_adult/ | https://bioconductor.org/packages/scRNAseq/ |
| LaMannoBrain\_mouse\_embryo | 1907 | scRNASeq | https://cola-rh.github.io/LaMannoBrain\_mouse\_embryo/ | https://bioconductor.org/packages/scRNAseq/ |
| LawlorPancreas | 638 | scRNASeq | https://cola-rh.github.io/LawlorPancreas/ | https://bioconductor.org/packages/scRNAseq/ |
| LengESC | 460 | scRNASeq | https://cola-rh.github.io/LengESC/ | https://bioconductor.org/packages/scRNAseq/ |
| ReprocessedTh2 | 96 | scRNASeq | https://cola-rh.github.io/ReprocessedTh2/ | https://bioconductor.org/packages/scRNAseq/ |
| MessmerESC | 1344 | scRNASeq | https://cola-rh.github.io/MessmerESC/ | https://bioconductor.org/packages/scRNAseq/ |
| MuraroPancreas | 3072 | scRNASeq | https://cola-rh.github.io/MuraroPancreas/ | https://bioconductor.org/packages/scRNAseq/ |
| NestorowaHSC | 1920 | scRNASeq | https://cola-rh.github.io/NestorowaHSC/ | https://bioconductor.org/packages/scRNAseq/ |
| PollenGlia | 367 | scRNASeq | https://cola-rh.github.io/PollenGlia/ | https://bioconductor.org/packages/scRNAseq/ |
| RichardTCell | 572 | scRNASeq | https://cola-rh.github.io/RichardTCell/ | https://bioconductor.org/packages/scRNAseq/ |
| RomanovBrain | 2881 | scRNASeq | https://cola-rh.github.io/RomanovBrain/ | https://bioconductor.org/packages/scRNAseq/ |
| SegerstolpePancreas | 3514 | scRNASeq | https://cola-rh.github.io/SegerstolpePancreas/ | https://bioconductor.org/packages/scRNAseq/ |
| UsoskinBrain | 864 | scRNASeq | https://cola-rh.github.io/UsoskinBrain/ | https://bioconductor.org/packages/scRNAseq/ |
| TasicBrain | 1809 | scRNASeq | https://cola-rh.github.io/TasicBrain/ | https://bioconductor.org/packages/scRNAseq/ |
| ReprocessedAllen | 379 | scRNASeq | https://cola-rh.github.io/ReprocessedAllen/ | https://bioconductor.org/packages/scRNAseq/ |
| XinPancreas | 1600 | scRNASeq | https://cola-rh.github.io/XinPancreas/ | https://bioconductor.org/packages/scRNAseq/ |
| ZeiselBrain | 3005 | scRNASeq | https://cola-rh.github.io/ZeiselBrain/ | https://bioconductor.org/packages/scRNAseq/ |
| Fluidigm | 65 | scRNASeq | https://cola-rh.github.io/Fluidigm/ | https://bioconductor.org/packages/scRNAseq/ |
| TCGA\_ACC\_methylation | 80 | 450K methylation array | https://cola-rh.github.io/TCGA\_ACC\_methylation/ | https://xenabrowser.net/datapages/?hub=https://tcga.xenahubs.net:443 |
| TCGA\_BLCA\_methylation | 434 | 450K methylation array | https://cola-rh.github.io/TCGA\_BLCA\_methylation/ | https://xenabrowser.net/datapages/?hub=https://tcga.xenahubs.net:443 |
| TCGA\_BRCA\_methylation | 888 | 450K methylation array | https://cola-rh.github.io/TCGA\_BRCA\_methylation/ | https://xenabrowser.net/datapages/?hub=https://tcga.xenahubs.net:443 |
| TCGA\_CESC\_methylation | 312 | 450K methylation array | https://cola-rh.github.io/TCGA\_CESC\_methylation/ | https://xenabrowser.net/datapages/?hub=https://tcga.xenahubs.net:443 |
| TCGA\_COAD\_methylation | 337 | 450K methylation array | https://cola-rh.github.io/TCGA\_COAD\_methylation/ | https://xenabrowser.net/datapages/?hub=https://tcga.xenahubs.net:443 |
| TCGA\_COADREAD\_methylation | 443 | 450K methylation array | https://cola-rh.github.io/TCGA\_COADREAD\_methylation/ | https://xenabrowser.net/datapages/?hub=https://tcga.xenahubs.net:443 |
| TCGA\_ESCA\_methylation | 202 | 450K methylation array | https://cola-rh.github.io/TCGA\_ESCA\_methylation/ | https://xenabrowser.net/datapages/?hub=https://tcga.xenahubs.net:443 |
| TCGA\_GBM\_methylation | 155 | 450K methylation array | https://cola-rh.github.io/TCGA\_GBM\_methylation/ | https://xenabrowser.net/datapages/?hub=https://tcga.xenahubs.net:443 |
| TCGA\_GBMLGG\_methylation | 685 | 450K methylation array | https://cola-rh.github.io/TCGA\_GBMLGG\_methylation/ | https://xenabrowser.net/datapages/?hub=https://tcga.xenahubs.net:443 |
| TCGA\_HNSC\_methylation | 580 | 450K methylation array | https://cola-rh.github.io/TCGA\_HNSC\_methylation/ | https://xenabrowser.net/datapages/?hub=https://tcga.xenahubs.net:443 |
| TCGA\_KICH\_methylation | 66 | 450K methylation array | https://cola-rh.github.io/TCGA\_KICH\_methylation/ | https://xenabrowser.net/datapages/?hub=https://tcga.xenahubs.net:443 |
| TCGA\_KIRC\_methylation | 480 | 450K methylation array | https://cola-rh.github.io/TCGA\_KIRC\_methylation/ | https://xenabrowser.net/datapages/?hub=https://tcga.xenahubs.net:443 |
| TCGA\_KIRP\_methylation | 321 | 450K methylation array | https://cola-rh.github.io/TCGA\_KIRP\_methylation/ | https://xenabrowser.net/datapages/?hub=https://tcga.xenahubs.net:443 |
| TCGA\_LAML\_methylation | 194 | 450K methylation array | https://cola-rh.github.io/TCGA\_LAML\_methylation/ | https://xenabrowser.net/datapages/?hub=https://tcga.xenahubs.net:443 |
| TCGA\_LGG\_methylation | 530 | 450K methylation array | https://cola-rh.github.io/TCGA\_LGG\_methylation/ | https://xenabrowser.net/datapages/?hub=https://tcga.xenahubs.net:443 |
| TCGA\_LIHC\_methylation | 429 | 450K methylation array | https://cola-rh.github.io/TCGA\_LIHC\_methylation/ | https://xenabrowser.net/datapages/?hub=https://tcga.xenahubs.net:443 |
| TCGA\_LUAD\_methylation | 492 | 450K methylation array | https://cola-rh.github.io/TCGA\_LUAD\_methylation/ | https://xenabrowser.net/datapages/?hub=https://tcga.xenahubs.net:443 |
| TCGA\_LUNG\_methylation | 907 | 450K methylation array | https://cola-rh.github.io/TCGA\_LUNG\_methylation/ | https://xenabrowser.net/datapages/?hub=https://tcga.xenahubs.net:443 |
| TCGA\_LUSC\_methylation | 415 | 450K methylation array | https://cola-rh.github.io/TCGA\_LUSC\_methylation/ | https://xenabrowser.net/datapages/?hub=https://tcga.xenahubs.net:443 |
| TCGA\_MESO\_methylation | 87 | 450K methylation array | https://cola-rh.github.io/TCGA\_MESO\_methylation/ | https://xenabrowser.net/datapages/?hub=https://tcga.xenahubs.net:443 |
| TCGA\_PAAD\_methylation | 195 | 450K methylation array | https://cola-rh.github.io/TCGA\_PAAD\_methylation/ | https://xenabrowser.net/datapages/?hub=https://tcga.xenahubs.net:443 |
| TCGA\_PCPG\_methylation | 187 | 450K methylation array | https://cola-rh.github.io/TCGA\_PCPG\_methylation/ | https://xenabrowser.net/datapages/?hub=https://tcga.xenahubs.net:443 |
| TCGA\_PRAD\_methylation | 549 | 450K methylation array | https://cola-rh.github.io/TCGA\_PRAD\_methylation/ | https://xenabrowser.net/datapages/?hub=https://tcga.xenahubs.net:443 |
| TCGA\_READ\_methylation | 106 | 450K methylation array | https://cola-rh.github.io/TCGA\_READ\_methylation/ | https://xenabrowser.net/datapages/?hub=https://tcga.xenahubs.net:443 |
| TCGA\_SARC\_methylation | 269 | 450K methylation array | https://cola-rh.github.io/TCGA\_SARC\_methylation/ | https://xenabrowser.net/datapages/?hub=https://tcga.xenahubs.net:443 |
| TCGA\_SKCM\_methylation | 476 | 450K methylation array | https://cola-rh.github.io/TCGA\_SKCM\_methylation/ | https://xenabrowser.net/datapages/?hub=https://tcga.xenahubs.net:443 |
| TCGA\_STAD\_methylation | 398 | 450K methylation array | https://cola-rh.github.io/TCGA\_STAD\_methylation/ | https://xenabrowser.net/datapages/?hub=https://tcga.xenahubs.net:443 |
| TCGA\_TGCT\_methylation | 156 | 450K methylation array | https://cola-rh.github.io/TCGA\_TGCT\_methylation/ | https://xenabrowser.net/datapages/?hub=https://tcga.xenahubs.net:443 |
| TCGA\_THCA\_methylation | 571 | 450K methylation array | https://cola-rh.github.io/TCGA\_THCA\_methylation/ | https://xenabrowser.net/datapages/?hub=https://tcga.xenahubs.net:443 |
| TCGA\_THYM\_methylation | 126 | 450K methylation array | https://cola-rh.github.io/TCGA\_THYM\_methylation/ | https://xenabrowser.net/datapages/?hub=https://tcga.xenahubs.net:443 |
| TCGA\_UCEC\_methylation | 478 | 450K methylation array | https://cola-rh.github.io/TCGA\_UCEC\_methylation/ | https://xenabrowser.net/datapages/?hub=https://tcga.xenahubs.net:443 |
| TCGA\_UCS\_methylation | 57 | 450K methylation array | https://cola-rh.github.io/TCGA\_UCS\_methylation/ | https://xenabrowser.net/datapages/?hub=https://tcga.xenahubs.net:443 |
| TCGA\_UVM\_methylation | 80 | 450K methylation array | https://cola-rh.github.io/TCGA\_UVM\_methylation/ | https://xenabrowser.net/datapages/?hub=https://tcga.xenahubs.net:443 |
