## Supplementary File 4 for "Improve consensus partitioning via a hierarchical procedure"

Compare cola HCP and Seurat classifications


### Compare cola HCP and Seurat classifications

###### Zuguang Gu

#### 2021-07-29

In Figure 5 of the manuscript, we compared the classifications from cola HCP and Seurat. In general, the two classifications are very similar, except the following differences:

- Group “01” in cola HCP is split into two groups with labels “0” and “2” in Seurat.
- Samples in cola HCP group “0212” and “022” have a different classification in Seurat with group labels “1” and “7”.

In this supplementary, we go deeper to see where are the differences between the two classifications.

Figure S4.1. Correspondance between cola HCP and Seurat classifications.

#### Compare cola HCP group “01” and Seurat group “0”/“2”

First we extract samples under cola HCP group “01” or samples under Seurat group “0”/“2”. The common samples in the two classifications are used for further analysis.

```
library(cola)
library(grid)
library(circlize)
library(ComplexHeatmap)
library(GetoptLong)

rh = readRDS("PBMC_cola_hierarchical_partition.rds")
load("Seurat_classification.RData")

mat = get_matrix(rh)

# group "01" in cola HCP classification
cl = rh@subgroup
l1 = cl %in% "01"
m1 = mat[, l1]
cl1 = get_classes(rh)[l1]

# group 0 and 2 in Seurat classification
names(Seurat_class) = colnames(mat)
l2 = Seurat_class %in% c(0, 2)
m2 = mat[, l2]
cl2 = Seurat_class[l2]

# the intersection of samples
ncol(m1)
```

```
## [1] 1193
```

```
ncol(m2)
```

```
## [1] 1183
```

```
cn = intersect(colnames(m1), colnames(m2))
length(cn)
```

```
## [1] 1139
```

The matrix of the subset of samples and the corresponding annotation table:

```
mm = mat[, cn]
anno = data.frame(cola_HCP = cl1[cn], Seurat_class = cl2[cn])

anno_col = list(
    cola_HCP = rh@subgroup_col[unique(cl[cn])],
    Seurat_class = Seurat_col[unique(cl2[cn])]
)
```

To compare the two classifications, one way is to compare the signature genes that are significantly expressed between the groups in the classification. For Seurat classification which contains two groups in the subset of samples, we apply *t*-test to look for significantly differentially expressed genes. FDRs are saved in the variable `fdr` and the *t*-values are saved in the variable `tvalue`.

```
library(genefilter)
fdr = list()
tvalue = list()
stat = rowttests(mm, factor(anno$Seurat_class))
fdr$Seurat_class = p.adjust(stat$p.value)
tvalue$Seurat_class = stat$statistic
fdr = as.data.frame(fdr)
tvalue = as.data.frame(tvalue)
```

cola HCP did not separate this subset of samples while marked them as a leaf node “01” in the hierarchy. We can check the consensus partitioning result at node “01”:

```
rh["01"]
```

```
## A 'DownSamplingConsensusPartition' object with k = 2, 3, 4.
##   On a matrix with 3664 rows and 500 columns, randomly sampled from 1193 columns.
##   Top rows (200) are extracted by 'ATC' method.
##   Subgroups are detected by 'skmeans' method.
##   Performed in total 150 partitions by row resampling.
##   Best k for subgroups seems to be 2.
## 
## Following methods can be applied to this 'DownSamplingConsensusPartition' object:
##  [1] "cola_report"             "collect_classes"        
##  [3] "collect_plots"           "collect_stats"          
##  [5] "colnames"                "compare_partitions"     
##  [7] "compare_signatures"      "consensus_heatmap"      
##  [9] "dimension_reduction"     "functional_enrichment"  
## [11] "get_anno"                "get_anno_col"           
## [13] "get_classes"             "get_consensus"          
## [15] "get_matrix"              "get_membership"         
## [17] "get_param"               "get_signatures"         
## [19] "get_stats"               "is_best_k"              
## [21] "is_stable_k"             "membership_heatmap"     
## [23] "ncol"                    "nrow"                   
## [25] "plot_ecdf"               "predict_classes"        
## [27] "rownames"                "select_partition_number"
## [29] "show"                    "suggest_best_k"         
## [31] "test_to_known_factors"   "top_rows_heatmap"
```

```
select_partition_number(rh["01"])
```

Figure S4.2 Various metrics to select the best number of subgroups for the consensus partitioning on node “01”.

The result shows the consensus partitioning result (k = 2) on node “01” is not “very stable” that it did not pass the default cutoff of silhouette scores (>= 0.95), thus, in HCP, this subset of samples was not split further more. (Readers can also try `collect_plots(rh["01"])` to get more diagnostic plots.)

We can still manually split node “01” into two subgroups and compare to the Seurat classification. In the following code, the 2-group cola CP classification on node “01” is added to the annotation table `anno`, and we also calcualte the differential expression for cola CP classification.

```
cl_CP = get_classes(rh["01"], k = 2)[cn, "class"]
anno$cola_CP = as.character(cl_CP)
stat = rowttests(mm, factor(anno$cola_CP))
fdr$cola_CP = p.adjust(stat$p.value)
tvalue$cola_CP = stat$statistic

anno_col$cola_CP = c("1" = "orange", "2" = "purple")
```

Next we visualize the differential genes (FDR < 0.05) in the two classifications.

Figure S4.3. Heatmap of signature genes from the two classifications.

And the overlap of the two sets of signature genes:

```
gl = list("cola_CP" = rownames(mm)[fdr$cola_CP < 0.05],
          "Seurat_class" = rownames(mm)[fdr$Seurat_class < 0.05])
library(eulerr)
plot(euler(gl), quantities = TRUE)
```

Figure S4.4. Euler diagram of the signature genes from the two classifications.

So, here in general, Seurat classification is similar to cola CP classification for this subset of samples. If we check the overlap of the two classifications:

```
table(anno$Seurat_class, anno$cola_CP)
```

```
##    
##       1   2
##   0 559 133
##   2  84 363
```

The similarity is:

```
(559 + 363)/nrow(anno)
```

```
## [1] 0.809482
```

But note, the cola CP classification is less stable by the sense of consensus partitioning.

The two classifications have similar sets of signature genes but the classifications are slightly different. We next check the differential expression in the two classifications by comparing the *t*-values from the *t*-tests.

```
plot(tvalue$Seurat_class, tvalue$cola_CP, asp = 1)
abline(a = 0, b = 1)
```

Figure S4.5. Compare differential expression in the two classifications.

The plot shows the differential expression in cola CP is higher than Seurat. For this sense, we can make the conclusion that cola can classify samples that are more separated than Seurat.

#### Compare cola HCP group “0212”/“022” with Seurat group “1”/“7”

Similarly, we compare the cola HCP classification with groups “0212”, “022” and the Seurat classification with groups “1” and “7”. We first extract the subset of samples.

```
# group "0212", "022" in cola HCP classification
cl = rh@subgroup
l1 = cl %in% c("0212", "022")
m1 = mat[, l1]
cl1 = get_classes(rh)[l1]

# group 1 and 7 in Seurat classification
names(Seurat_class) = colnames(rh)
l2 = Seurat_class %in% c(1, 7)
m2 = mat[, l2]
cl2 = Seurat_class[l2]

# the intersection of samples
ncol(m1)
```

```
## [1] 529
```

```
ncol(m2)
```

```
## [1] 512
```

```
cn = intersect(colnames(m1), colnames(m2))
length(cn)
```

```
## [1] 505
```

```
# the submatrix
mm = mat[, cn]
anno = data.frame(cola_HCP = cl1[cn], Seurat_class = cl2[cn])

anno_col = list(
    cola_HCP = rh@subgroup_col[unique(cl[cn])],
    Seurat_class = Seurat_col[unique(cl2[cn])]
)
```

We apply differential expression analysis to both cola HCP classification and Seurat classification.

```
library(genefilter)
fdr = list()
tvalue = list()
stat = rowttests(mm, factor(anno$Seurat_class))
fdr$Seurat_class = p.adjust(stat$p.value)
tvalue$Seurat_class = stat$statistic

stat = rowttests(mm, factor(anno$cola_HCP))
fdr$cola_HCP = p.adjust(stat$p.value)
tvalue$cola_HCP = stat$statistic

fdr = as.data.frame(fdr)
tvalue = as.data.frame(tvalue)
```

Heatmaps of the two sets of signature genes.

Figure S4.6. Heatmap of signature genes from the two classifications.

The overlap of the two sets of signature genes. Here we can see the two sets of signatures are quite different.

```
gl = list("cola_HCP" = rownames(mm)[fdr$cola_HCP < 0.05],
          "Seurat_class" = rownames(mm)[fdr$Seurat_class < 0.05])
library(eulerr)
plot(euler(gl), quantities = TRUE)
```

Figure S4.7. Euler diagram of the signature genes from the two classifications.

Then it is worthwhile to compare the bioloigcal functions (by Gene Ontology terms) of the two different sets of signature genes.

```
fl = lapply(gl, functional_enrichment)
library(simplifyEnrichment)
simplifyGOFromMultipleLists(lapply(fl, function(x) x$BP))
```

Figure S4.8. Functional enrichment analysis on the signature gene lists.

Generally, both lists of signature genes generate quite a lot of significant GO terms and their biological functions are similar. cola HCP generates more significant GO terms (208) than Seurat (132) under FDR < 0.01.

#### Session info

```
sessionInfo()
```

```
## R version 4.1.0 (2021-05-18)
## Platform: x86_64-apple-darwin17.0 (64-bit)
## Running under: macOS Big Sur 10.16
## 
## Matrix products: default
## BLAS:   /Library/Frameworks/R.framework/Versions/4.1/Resources/lib/libRblas.dylib
## LAPACK: /Library/Frameworks/R.framework/Versions/4.1/Resources/lib/libRlapack.dylib
## 
## locale:
## [1] C/UTF-8/C/C/C/C
## 
## attached base packages:
##  [1] parallel  stats4    grid      stats     graphics  grDevices utils    
##  [8] datasets  methods   base     
## 
## other attached packages:
##  [1] simplifyEnrichment_1.3.1 org.Hs.eg.db_3.13.0      AnnotationDbi_1.54.1    
##  [4] IRanges_2.26.0           S4Vectors_0.30.0         Biobase_2.52.0          
##  [7] BiocGenerics_0.38.0      eulerr_6.1.0             cowplot_1.1.1           
## [10] genefilter_1.74.0        GetoptLong_1.0.5         ComplexHeatmap_2.9.3    
## [13] circlize_0.4.13          cola_1.9.4               knitr_1.33              
## [16] rmarkdown_2.9            BiocManager_1.30.16      colorout_1.2-2          
## 
## loaded via a namespace (and not attached):
##   [1] shadowtext_0.0.8       fastmatch_1.1-0        plyr_1.8.6            
##   [4] igraph_1.2.6           lazyeval_0.2.2         proxyC_0.2.0          
##   [7] polylabelr_0.2.0       splines_4.1.0          BiocParallel_1.26.0   
##  [10] GenomeInfoDb_1.28.0    ggplot2_3.3.5          digest_0.6.27         
##  [13] foreach_1.5.1          htmltools_0.5.1.1      GOSemSim_2.18.0       
##  [16] viridis_0.6.1          magick_2.7.2           GO.db_3.13.0          
##  [19] fansi_0.5.0            magrittr_2.0.1         memoise_2.0.0         
##  [22] tm_0.7-8               cluster_2.1.2          doParallel_1.0.16     
##  [25] Biostrings_2.60.1      annotate_1.70.0        graphlayouts_0.7.1    
##  [28] RcppParallel_5.1.4     matrixStats_0.59.0     enrichplot_1.12.1     
##  [31] colorspace_2.0-2       blob_1.2.1             ggrepel_0.9.1         
##  [34] xfun_0.24              dplyr_1.0.7            crayon_1.4.1          
##  [37] RCurl_1.98-1.3         microbenchmark_1.4-7   jsonlite_1.7.2        
##  [40] scatterpie_0.1.6       impute_1.66.0          ape_5.5               
##  [43] brew_1.0-6             survival_3.2-11        iterators_1.0.13      
##  [46] glue_1.4.2             polyclip_1.10-0        gtable_0.3.0          
##  [49] zlibbioc_1.38.0        XVector_0.32.0         shape_1.4.6           
##  [52] scales_1.1.1           DOSE_3.18.1            bezier_1.1.2          
##  [55] DBI_1.1.1              Rcpp_1.0.6             gridtext_0.1.4        
##  [58] viridisLite_0.4.0      xtable_1.8-4           clue_0.3-59           
##  [61] tidytree_0.3.4         bit_4.0.4              mclust_5.4.7          
##  [64] httr_1.4.2             fgsea_1.18.0           RColorBrewer_1.1-2    
##  [67] ellipsis_0.3.2         pkgconfig_2.0.3        XML_3.99-0.6          
##  [70] farver_2.1.0           sass_0.4.0             utf8_1.2.1            
##  [73] tidyselect_1.1.1       rlang_0.4.11           reshape2_1.4.4        
##  [76] munsell_0.5.0          tools_4.1.0            cachem_1.0.5          
##  [79] downloader_0.4         generics_0.1.0         RSQLite_2.2.7         
##  [82] evaluate_0.14          stringr_1.4.0          fastmap_1.1.0         
##  [85] yaml_2.2.1             ggtree_3.0.2           bit64_4.0.5           
##  [88] tidygraph_1.2.0        purrr_0.3.4            KEGGREST_1.32.0       
##  [91] ggraph_2.0.5           nlme_3.1-152           slam_0.1-48           
##  [94] aplot_0.0.6            DO.db_2.9              xml2_1.3.2            
##  [97] compiler_4.1.0         png_0.1-7              treeio_1.16.1         
## [100] tibble_3.1.2           tweenr_1.0.2           bslib_0.2.5.1         
## [103] stringi_1.6.2          highr_0.9              lattice_0.20-44       
## [106] Matrix_1.3-4           markdown_1.1           vctrs_0.3.8           
## [109] pillar_1.6.1           lifecycle_1.0.0        jquerylib_0.1.4       
## [112] GlobalOptions_0.1.2    data.table_1.14.0      bitops_1.0-7          
## [115] irlba_2.3.3            patchwork_1.1.1        qvalue_2.24.0         
## [118] R6_2.5.0               gridExtra_2.3          codetools_0.2-18      
## [121] MASS_7.3-54            assertthat_0.2.1       rjson_0.2.20          
## [124] GenomeInfoDbData_1.2.6 clusterProfiler_4.0.0  tidyr_1.1.3           
## [127] rvcheck_0.1.8          skmeans_0.2-13         Cairo_1.5-12.2        
## [130] ggforce_0.3.3          NLP_0.2-1
```
