## Supplementary File 5 for "Improve consensus partitioning via a hierarchical procedure"

Classification of central nervous system tumors


### Classification of central nervous system tumors

###### Zuguang Gu

#### 2021-09-03

Capper et. al., 2018 studied subtype classification of central nervous system tumors from DNA methylation data. In their dataset, there are 14 different tumor types (include controls) which are classified into 91 subtypes inferred from 2801 samples.

```
library(spiralize)
library(ComplexHeatmap)
df = readRDS(system.file("extdata", "CNS_tumour_classification.rds", package = "spiralize"))
n = nrow(df)
spiral_rle = function(x, col, labels = FALSE) {
    x = as.vector(x) # in case it is a factor
    r1 = rle(x)
    for(i in seq_along(r1$lengths)) {
        spiral_rect(sum(r1$lengths[seq_len(i-1)]), 0, sum(r1$lengths[seq_len(i)]), 1, gp = gpar(fill = col[r1$values[i]], col = NA))
    }

    if(labels) {
        for(i in seq_along(r1$lengths)) {
            spiral_text( (sum(r1$lengths[seq_len(i-1)]) + sum(r1$lengths[seq_len(i)]))/2, 0.5, r1$values[i], 
                facing = "curved_inside", nice_facing = TRUE)
        }
    }
}

spiral_initialize(xlim = c(0, n), scale_by = "curve_length", 
    vp_param = list(x = unit(0, "npc"), just = "left"))
spiral_track(height = 0.4)
meth_col = structure(names = unique(df$meth_class), unique(df$meth_col))
spiral_rle(df$meth_class, col = meth_col)

spiral_track(height = 0.4)
tumor_col = structure(names = unique(as.vector(df$tumor_type)), unique(df$tumor_col))
spiral_rle(df$tumor_type, col = tumor_col, labels = TRUE)

lgd_list = tapply(1:nrow(df), df$tumor_type, function(ind) {
    Legend(title = df$tumor_type[ind][1], at = unique(df$meth_class[ind]),
        legend_gp = gpar(fill = unique(df$meth_col[ind])))
})

# here set max_height to the height of the image so that the legends are automatically arranged
lgd = packLegend(list = lgd_list, max_height = unit(7, "inch"))
draw(lgd, x = unit(1, "npc") + unit(2, "mm"), just = "left")
```

Figure S5.1. Classification of central nervous system tumors.

#### Session info

```
sessionInfo()
```

```
## R version 4.1.0 (2021-05-18)
## Platform: x86_64-apple-darwin17.0 (64-bit)
## Running under: macOS Big Sur 10.16
## 
## Matrix products: default
## BLAS:   /Library/Frameworks/R.framework/Versions/4.1/Resources/lib/libRblas.dylib
## LAPACK: /Library/Frameworks/R.framework/Versions/4.1/Resources/lib/libRlapack.dylib
## 
## locale:
## [1] C/UTF-8/C/C/C/C
## 
## attached base packages:
## [1] grid      stats     graphics  grDevices utils     datasets  methods  
## [8] base     
## 
## other attached packages:
## [1] ComplexHeatmap_2.9.3  spiralize_1.0.2       knitr_1.33           
## [4] rmarkdown_2.9         preprocessCore_1.54.0 RColorBrewer_1.1-2   
## [7] cola_1.9.4            BiocManager_1.30.16   colorout_1.2-2       
## 
## loaded via a namespace (and not attached):
##   [1] colorspace_2.0-2       rjson_0.2.20           ellipsis_0.3.2        
##   [4] mclust_5.4.7           circlize_0.4.13        markdown_1.1          
##   [7] XVector_0.32.0         GlobalOptions_0.1.2    clue_0.3-59           
##  [10] rstudioapi_0.13        bit64_4.0.5            lubridate_1.7.10      
##  [13] AnnotationDbi_1.54.1   Polychrome_1.2.6       fansi_0.5.0           
##  [16] xml2_1.3.2             codetools_0.2-18       splines_4.1.0         
##  [19] doParallel_1.0.16      cachem_1.0.5           impute_1.66.0         
##  [22] jsonlite_1.7.2         Cairo_1.5-12.2         annotate_1.70.0       
##  [25] cluster_2.1.2          png_0.1-7              data.tree_1.0.0       
##  [28] compiler_4.1.0         httr_1.4.2             assertthat_0.2.1      
##  [31] Matrix_1.3-4           fastmap_1.1.0          htmltools_0.5.1.1     
##  [34] tools_4.1.0            gtable_0.3.0           glue_1.4.2            
##  [37] GenomeInfoDbData_1.2.6 dplyr_1.0.7            Rcpp_1.0.7            
##  [40] slam_0.1-48            Biobase_2.52.0         eulerr_6.1.0          
##  [43] jquerylib_0.1.4        vctrs_0.3.8            Biostrings_2.60.1     
##  [46] iterators_1.0.13       xfun_0.24              stringr_1.4.0         
##  [49] lifecycle_1.0.0        irlba_2.3.3            XML_3.99-0.6          
##  [52] dendextend_1.15.1      zlibbioc_1.38.0        scales_1.1.1          
##  [55] microbenchmark_1.4-7   parallel_4.1.0         yaml_2.2.1            
##  [58] memoise_2.0.0          gridExtra_2.3          ggplot2_3.3.5         
##  [61] sass_0.4.0             stringi_1.6.2          RSQLite_2.2.7         
##  [64] highr_0.9              genefilter_1.74.0      S4Vectors_0.30.0      
##  [67] foreach_1.5.1          BiocGenerics_0.38.0    shape_1.4.6           
##  [70] GenomeInfoDb_1.28.0    rlang_0.4.11           pkgconfig_2.0.3       
##  [73] matrixStats_0.59.0     bitops_1.0-7           evaluate_0.14         
##  [76] lattice_0.20-44        purrr_0.3.4            bit_4.0.4             
##  [79] tidyselect_1.1.1       magrittr_2.0.1         R6_2.5.0              
##  [82] IRanges_2.26.0         magick_2.7.2           generics_0.1.0        
##  [85] DBI_1.1.1              pillar_1.6.1           survival_3.2-11       
##  [88] KEGGREST_1.32.0        scatterplot3d_0.3-41   RCurl_1.98-1.3        
##  [91] tibble_3.1.2           crayon_1.4.1           utf8_1.2.1            
##  [94] skmeans_0.2-13         viridis_0.6.1          GetoptLong_1.0.3      
##  [97] blob_1.2.1             digest_0.6.27          xtable_1.8-4          
## [100] brew_1.0-6             stats4_4.1.0           munsell_0.5.0         
## [103] viridisLite_0.4.0      bslib_0.2.5.1
```
