## Supplementary File 6 for "Improve consensus partitioning via a hierarchical procedure"

Analysis of the TCGA GBM microarray dataset


### Analysis of the TCGA GBM microarray dataset

###### Zuguang Gu

#### 2021-09-03

We read the matrix and perform normalization.

```
library(cola)

m = read.table("https://jokergoo.github.io/cola_examples/TCGA_GBM/unifiedScaled.txt", 
    header = TRUE, row.names = 1, check.names = FALSE)
m = as.matrix(m)

subtype = read.table("https://jokergoo.github.io/cola_examples/TCGA_GBM/TCGA_unified_CORE_ClaNC840.txt", 
    sep = "\t", header = TRUE, check.names = FALSE, stringsAsFactors = FALSE)
subtype = structure(unlist(subtype[1, -(1:2)]), names = colnames(subtype)[-(1:2)])
subtype_col = structure(seq_len(4), names = unique(subtype))

m = m[, names(subtype)]
m = adjust_matrix(m)

library(preprocessCore)
cn = colnames(m)
rn = rownames(m)
m = normalize.quantiles(m)
colnames(m) = cn
rownames(m) = rn
```

First we apply standard consensus partitioning analysis with “ATC” as the top-value method and “skmeans” as partitioning method.

```
set.seed(123)
res = consensus_partition(m, top_value_method = "ATC", partition_method = "skmeans",
    cores = 4, anno = subtype, anno_col = subtype_col)
```

In the following plot, **cola** suggests 5 as the best number of subgroups, but we select 4 as the best k because it gives more stable classification.

```
suggest_best_k(res)
```

```
## The best k suggested by this function might not reflect the real
## subgroups in the data (especially when you expect a large best k). It
## is recommended to directly look at the plots from
## select_partition_number() or other related plotting functions.
```

```
## [1] 5
## attr(,"optional")
## [1] 2 3 4
```

```
select_partition_number(res)
```

Figure S6.1. Select the best number of groups.

The signature heatmap with 4 subgroups.

```
get_signatures(res, k = 4)
```

Figure S6.2. Signature heatmap of CP classification with 4 subgroups.

Next we apply hierarchical consensus partitioning (HCP) on the same matrix:

```
set.seed(123)
rh = hierarchical_partition(m, cores = 4, anno = subtype, anno_col = subtype_col)
```

The subgroup hierarchy:

```
collect_classes(rh)
```

Figure S6.3. Subgroup hierarchy under HCP.

And the signature heatmap under HCP classification:

```
get_signatures(rh)
```

Figure S6.4. Signature heatmap under HCP classification.

The statistics on each node:

```
df = node_info(rh)
df
```

```
##      id best_method depth best_k n_columns n_signatures p_signatures is_leaf
## 1     0 ATC:skmeans     1      4       173         9686 0.8596787077   FALSE
## 2    01 ATC:skmeans     2      3        52         4051 0.3595455756   FALSE
## 3   011 ATC:skmeans     3      2        17           55 0.0048815124    TRUE
## 4   012 ATC:skmeans     3      2        21          625 0.0554717316   FALSE
## 5  0121 not applied     4     NA        10           NA           NA    TRUE
## 6  0122 not applied     4     NA        11           NA           NA    TRUE
## 7   013 ATC:skmeans     3      2        14            8 0.0007100382    TRUE
## 8    02 ATC:skmeans     2      3        66         4781 0.4243365581   FALSE
## 9   021 ATC:skmeans     3      3        25          806 0.0715363451   FALSE
## 10 0211 not applied     4     NA        11           NA           NA    TRUE
## 11 0212 not applied     4     NA         9           NA           NA    TRUE
## 12 0213 not applied     4     NA         5           NA           NA    TRUE
## 13  022 ATC:skmeans     3      2        24          666 0.0591106772   FALSE
## 14 0221 ATC:skmeans     4      2        14           30 0.0026626431    TRUE
## 15 0222 not applied     4     NA        10           NA           NA    TRUE
## 16  023 ATC:skmeans     3      2        17          227 0.0201473329    TRUE
## 17   03 ATC:skmeans     2      4        24         1376 0.1221265643   FALSE
## 18  031 not applied     3     NA         7           NA           NA    TRUE
## 19  032 not applied     3     NA         6           NA           NA    TRUE
## 20  033 not applied     3     NA         6           NA           NA    TRUE
## 21  034 not applied     3     NA         5           NA           NA    TRUE
## 22   04 ATC:skmeans     2      2        31         1954 0.1734268217   FALSE
## 23  041 ATC:skmeans     3      2        18          257 0.0228099760    TRUE
## 24  042 ATC:skmeans     3      2        13          175 0.0155320848    TRUE
```

And the statistics on non-leaf nodes:

```
df[!df$is_leaf, ]
```

```
##     id best_method depth best_k n_columns n_signatures p_signatures is_leaf
## 1    0 ATC:skmeans     1      4       173         9686   0.85967871   FALSE
## 2   01 ATC:skmeans     2      3        52         4051   0.35954558   FALSE
## 4  012 ATC:skmeans     3      2        21          625   0.05547173   FALSE
## 8   02 ATC:skmeans     2      3        66         4781   0.42433656   FALSE
## 9  021 ATC:skmeans     3      3        25          806   0.07153635   FALSE
## 13 022 ATC:skmeans     3      2        24          666   0.05911068   FALSE
## 17  03 ATC:skmeans     2      4        24         1376   0.12212656   FALSE
## 22  04 ATC:skmeans     2      2        31         1954   0.17342682   FALSE
```

#### Session Info

```
sessionInfo()
```

```
## R version 4.1.0 (2021-05-18)
## Platform: x86_64-apple-darwin17.0 (64-bit)
## Running under: macOS Big Sur 10.16
## 
## Matrix products: default
## BLAS:   /Library/Frameworks/R.framework/Versions/4.1/Resources/lib/libRblas.dylib
## LAPACK: /Library/Frameworks/R.framework/Versions/4.1/Resources/lib/libRlapack.dylib
## 
## locale:
## [1] C/UTF-8/C/C/C/C
## 
## attached base packages:
## [1] stats     graphics  grDevices utils     datasets  methods   base     
## 
## other attached packages:
## [1] knitr_1.33            rmarkdown_2.9         preprocessCore_1.54.0
## [4] RColorBrewer_1.1-2    cola_1.9.4            BiocManager_1.30.16  
## [7] colorout_1.2-2       
## 
## loaded via a namespace (and not attached):
##   [1] colorspace_2.0-2       rjson_0.2.20           ellipsis_0.3.2        
##   [4] mclust_5.4.7           circlize_0.4.13        markdown_1.1          
##   [7] XVector_0.32.0         GlobalOptions_0.1.2    clue_0.3-59           
##  [10] rstudioapi_0.13        bit64_4.0.5            AnnotationDbi_1.54.1  
##  [13] Polychrome_1.2.6       fansi_0.5.0            xml2_1.3.2            
##  [16] codetools_0.2-18       splines_4.1.0          doParallel_1.0.16     
##  [19] cachem_1.0.5           impute_1.66.0          jsonlite_1.7.2        
##  [22] Cairo_1.5-12.2         annotate_1.70.0        cluster_2.1.2         
##  [25] png_0.1-7              data.tree_1.0.0        compiler_4.1.0        
##  [28] httr_1.4.2             assertthat_0.2.1       Matrix_1.3-4          
##  [31] fastmap_1.1.0          htmltools_0.5.1.1      tools_4.1.0           
##  [34] gtable_0.3.0           glue_1.4.2             GenomeInfoDbData_1.2.6
##  [37] dplyr_1.0.7            Rcpp_1.0.7             slam_0.1-48           
##  [40] Biobase_2.52.0         eulerr_6.1.0           jquerylib_0.1.4       
##  [43] vctrs_0.3.8            Biostrings_2.60.1      iterators_1.0.13      
##  [46] xfun_0.24              stringr_1.4.0          lifecycle_1.0.0       
##  [49] irlba_2.3.3            XML_3.99-0.6           dendextend_1.15.1     
##  [52] zlibbioc_1.38.0        scales_1.1.1           microbenchmark_1.4-7  
##  [55] parallel_4.1.0         ComplexHeatmap_2.9.3   yaml_2.2.1            
##  [58] memoise_2.0.0          gridExtra_2.3          ggplot2_3.3.5         
##  [61] sass_0.4.0             stringi_1.6.2          RSQLite_2.2.7         
##  [64] highr_0.9              genefilter_1.74.0      S4Vectors_0.30.0      
##  [67] foreach_1.5.1          BiocGenerics_0.38.0    shape_1.4.6           
##  [70] GenomeInfoDb_1.28.0    rlang_0.4.11           pkgconfig_2.0.3       
##  [73] matrixStats_0.59.0     bitops_1.0-7           evaluate_0.14         
##  [76] lattice_0.20-44        purrr_0.3.4            bit_4.0.4             
##  [79] tidyselect_1.1.1       magrittr_2.0.1         R6_2.5.0              
##  [82] IRanges_2.26.0         magick_2.7.2           generics_0.1.0        
##  [85] DBI_1.1.1              pillar_1.6.1           survival_3.2-11       
##  [88] KEGGREST_1.32.0        scatterplot3d_0.3-41   RCurl_1.98-1.3        
##  [91] tibble_3.1.2           crayon_1.4.1           utf8_1.2.1            
##  [94] skmeans_0.2-13         viridis_0.6.1          GetoptLong_1.0.3      
##  [97] grid_4.1.0             blob_1.2.1             digest_0.6.27         
## [100] xtable_1.8-4           brew_1.0-6             stats4_4.1.0          
## [103] munsell_0.5.0          viridisLite_0.4.0      bslib_0.2.5.1
```
